# Extreme allelic heterogeneity at a *Caenorhabditis elegans* beta-tubulin locus explains natural resistance to benzimidazoles

**DOI:** 10.1101/372623

**Authors:** Steffen R. Hahnel, Stefan Zdraljevic, Briana C. Rodriguez, Yuehui Zhao, Patrick T. McGrath, Erik C. Andersen

**Author notes:** Equal contribution (Steffen R. Hahnel, Stefan Zdraljevic). Corresponding author: **Erik C. Andersen**, Assistant Professor of Molecular Biosciences, Northwestern University, Evanston, IL 60208, USA. Steffen R. Hahnel, ORCID 0000-0001-8848-0691. Stefan Zdraljevic, ORCID 0000-0003-2883-4616. Patrick T. McGrath, ORCID 0000-0002-1598-3746. Erik C. Andersen, ORCID 0000-0003-0229-9651.

## Abstract

Benzimidazoles (BZ) are essential components of the limited chemotherapeutic arsenal available to control the global burden of parasitic nematodes. The emerging threat of BZ resistance among nearly all nematode species necessitates the development of novel strategies to identify genetic and molecular mechanisms underlying this resistance. All detection of parasitic helminth resistance to BZ is focused on the genotyping of three variant sites in the orthologs of the β-tubulin gene found to confer resistance in the free-living nematode *Caenorhabditis elegans*. Because of the limitations of laboratory and field experiments in parasitic nematodes, it is difficult to look beyond these three sites, and additional BZ resistance is observed in the field. Here, we took an unbiased genome-wide mapping approach in the free-living nematode species *C. elegans* to identify the genetic underpinnings of natural resistance to the commonly used BZ, albendazole (ABZ). We found a wide range of natural variation in ABZ resistance in natural *C. elegans* populations. In agreement with known mechanisms of BZ resistance in parasites, we find that a majority of the variation in ABZ resistance among wild *C. elegans* strains is caused by variation in the β-tubulin gene *ben-1*. This result shows empirically that resistance to ABZ naturally exists and segregates within the *C. elegans* population, suggesting that selection in natural niches could enrich for resistant alleles. We identified 25 distinct *ben-1* alleles that are segregating at low frequencies within the *C. elegans* population, including many novel molecular variants. Population genetic analyses indicate that *ben-1* variation arose multiple times during the evolutionary history of *C. elegans* and provide evidence that these alleles likely occurred recently because of local selective pressures. Additionally, we find purifying selection at all five β-tubulin genes, despite predicted loss-of-function resistants variants in *ben-1*, indicating that BZ resistance in natural niches is a stronger selective pressure than loss of one β-tubulin gene. Furthermore, we use genome-editing to show that the most common parasitic nematode β-tubulin allele that confers BZ resistance, F200Y, confers resistance in *C. elegans*. Importantly, we identified a novel genomic region that is correlated with ABZ resistance in the *C. elegans* population but independent of *ben-1* and the other β-tubulin loci, suggesting that there are multiple mechanisms underlying BZ resistance. Taken together, our results establish a population-level resource of nematode natural diversity as an important model for the study of mechanisms that give rise to BZ resistance.

**Author summary:** Nematode parasites have a tremendous impact on human health with almost two billion people infected worldwide. The control of nematode infections relies mainly on the efficacy of a limited repertoire of anthelmintic compounds, including the benzimidazoles (BZ). Already a significant problem in veterinary medicine, increasing evidence exists for the development of BZ resistance in nematodes that infect humans. Laboratory screens and field surveys identified β-tubulin genes as major determinants of BZ resistance in nematodes but detailed population-wide genetic analyses of resistance mechanisms are only just beginning. Therefore, we took advantage of the free-living model organism *Caenorhabditis elegans* to study the genetic basis of resistance to the commonly used BZ, albendazole (ABZ) in a natural nematode population. Performing genome-wide association mappings, we were able to identify extreme heterogeneity in the β-tubulin gene *ben-1* as a major determinant of ABZ resistance. Moreover, our study provided new insights into the effects of missense and loss-of-function alleles at this locus, and how anthelmintic resistance could have developed within a natural nematode population.

## Introduction

Parasitic nematodes have a tremendous impact on global health and socio-economic development, especially in the developing world [1]. They are among the most widespread human pathogens, and almost two billion people are estimated to suffer from infection of one or multiple nematode species [1,2]. The main endemic areas of nematode infections are highly correlated with tropical and subtropical regions worldwide. Because of their detrimental impact on human health, several nematode infections belong to a class of diseases designated by the World Health Organization (WHO) as Neglected Tropical Diseases (NTDs). Altogether, the loss of disability-adjusted life years (DALY) caused by parasitic nematodes is conservatively estimated to be 10 million DALYs per year, which ranks them among the top of all NTDs [1]. Apart from this drastic impact on human health, several nematode species infect a variety of key crops and livestock causing substantial economic losses throughout the world [3].

Global control of nematode infections relies on the efficacy of a limited repertoire of anthelmintic drugs, including benzimidazoles (BZ). BZs are frequently used in mass drug administration (MDA) programs to treat parasitic nematode infections in endemic regions. However, the long-term success of these MDA programs is limited by the persistence and re-emergence of nematode infections in these regions. The most significant of these factors is the high reinfection rate caused by long-term parasitic nematode reservoirs, which necessitates frequent deworming of affected communities [2]. As a consequence of continuous drug-pressure and relaxation cycles, parasite populations have the potential to develop anthelmintic resistance. This resistance is already a significant problem in veterinary medicine [4], and resistance in the parasitic nematodes that infect humans is becoming increasingly evident. For example, reduced cure rates have been observed for soil-transmitted helminthiases, which might be attributable to BZ resistance [5–7]. To face this growing threat, detailed knowledge of anthelmintic resistance mechanisms is required to improve diagnostic tools for field surveys and educate drug-treatment strategies in MDA programs.

*In vitro* studies have shown that BZs inhibit the polymerization of microtubules [8–10]. Mutagenesis screens that selected for BZ resistance in *Saccharomyces cerevisiae* [11] and *Caenorhabditis elegans* [12] have independently identified numerous β-tubulin mutant alleles. These findings were then translated to veterinary-relevant nematodes, where three major single-nucleotide variants (SNVs) in parasitic nematode β-tubulin genes were found to be highly correlated with BZ resistance. The most common variant causes a tyrosine to phenylalanine amino acid change at position 200 (F200Y) [13]. Other SNVs linked to BZ resistance have been described at positions 167 (F167Y) and 198 (E198A) in several nematode species [14,15]. Although all three of these missense variants have been shown to reduce BZ binding affinity to purified β-tubulins *in vitro* [16,17], the BZ β-tubulin binding site remains experimentally uncharacterized. Furthermore, these missense variants do not explain all of the BZ resistance observed in the field [4,18]. This discrepancy might indicate that additional, non-target related components such as drug efflux pumps and detoxification enzymes might contribute to BZ resistance [4,19,20].

Additional genetic variants associated with natural BZ resistance can be identified using quantitative genetic approaches that consider genotypic and phenotypic variation present within a wild population of parasitic nematodes. However, parasitic nematodes are not easily amenable to quantitative genetic approaches because of their complicated life cycles, their poorly annotated reference genomes, and limited molecular and genetic tools [21–23]. Recent successes in the hookworm parasites found genomic intervals that underlie ivermectin resistance [21,24], but similar approaches have not been applied to BZ resistance yet. By contrast, the free-living nematode species *C. elegans* has a short life cycle, a well annotated reference genome [25–27], and an abundance of molecular and genetic tools for the characterization of BZ responses [28–30]. A recent example that applied quantitative genetic approaches to investigated BZ responses in *C. elegans* used a mapping population generated between two genetically divergent strains to identify a quantitative trait loci (QTL) linked to BZ resistance [30]. Importantly, the QTL identified in this study did not overlap with β-tubulin genes, suggesting that quantitative genetic approaches in *C. elegans* can be used to discover novel mechanisms associated with BZ resistance.

In the present study, we leveraged the power of *C. elegans* natural diversity to perform genome-wide association (GWA) mappings for BZ resistance. We used a set of 249 wild *C. elegans* isolates available through the *C. elegans* Natural Diversity Resource (CeNDR) [27] to identify genomic regions that contribute to albendazole (ABZ) resistance. ABZ is a broadly administered BZ used to treat parasitic nematode infections in humans and livestock [4,30]. We show that a major source of ABZ resistance in the *C. elegans* population is driven by putative loss-of-function (LoF) variants in the *ben-1* locus. Notably, we found 25 distinct *ben-1* alleles with low minor allele frequencies (MAF) that contribute to ABZ resistance. We show that these putative *ben-1* LoF variants arose independently during the evolutionary history of the *C. elegans* species, which suggests that local BZ selective pressures might have contributed to the extreme allelic heterogeneity at this locus. We next made use of the extensive molecular toolkit available in *C. elegans* to verify that the introduction of the *ben-1* LoF alleles do not result in any detectable fitness consequences in standard laboratory conditions, but does confer ABZ resistance. Taken together, these results suggest that *C. elegans* and likely free-living stages of parasitic species encounter natural compounds that promote BZ resistance through selection for standing or *de novo* variation in conserved nematode-specific β-tubulin genes.

## Results

### Genetically distinct *C. elegans* natural isolates respond differently to ABZ

We investigated the differences in responses across a panel of *C. elegans* wild strains to one of the most commonly used BZ compounds (albendazole, ABZ) in human and veterinary medicine [4,27,37]. To test the effects of ABZ treatment on *C. elegans*, we exposed four genetically divergent *C. elegans* isolates to various ABZ concentrations and measured the number and length of progeny the animals produced. We used these two traits because they represent endpoint measures for ABZ efficacy against human parasitic nematodes. After four days of exposure to ABZ, we detected a decrease in brood size and progeny length for all four assayed strains (Supplemental Figure 1; Supplemental data 1 and 2). Additionally, this assay revealed differential ABZ sensitivity among these wild strains, as measured by brood size and progeny length. For subsequent genome-wide association (GWA) mapping experiments, we used an ABZ concentration of 12.5 μM, which is a concentration that induced robust differences in ABZ responses among the four assayed wild strains.

### Natural variation in *C. elegans* ABZ responses maps to multiple genomic regions, including the *ben-1* locus

To identify genomic loci that underlie strain-specific ABZ responses, we performed genome-wide association (GWA) mappings using HTA fitness data obtained from exposing 209 *C. elegans* to either DMSO (control) or ABZ and DMSO conditions (Supplemental data 3 and 4). We employed two statistical methods to perform GWA mappings using the regressed animal length (q90.TOF) and normalized brood size (norm.n) traits, a single-marker and a gene-burden approach [39,40]. Using the single-marker approach [32, 35–37], we identified three distinct quantitative trait loci (QTL) that explained variation in animal length among wild isolates exposed to ABZ, whereas the brood-size trait did not map to any significant genomic loci (Supplemental data 5 and 6). Two of the animal-length QTL we identified are located on chromosome II, and the third is located on chromosome V (Figure 1A). The QTL on the left arm of chromosome II spans from 25 kb to 3.9 Mb and has a peak-marker position at 458 kb. The second QTL on chromosome II has a peak-marker position at 11 Mb and spans from 10.17 Mb to 11.6 Mb. The QTL on the right arm of chromosome V spans from from 18.04 Mb to 18.68 Mb, with the peak marker located at 18.35 Mb (Supplemental Figure 2; Supplemental data 5). Remarkably, the β-tubulin gene *ben-1*, shown to be the major determinant for ABZ resistance in the *C. elegans* laboratory strain N2 [12,13], did not map using this single marker-based approach (Figure 1A).

**Figure 1:**
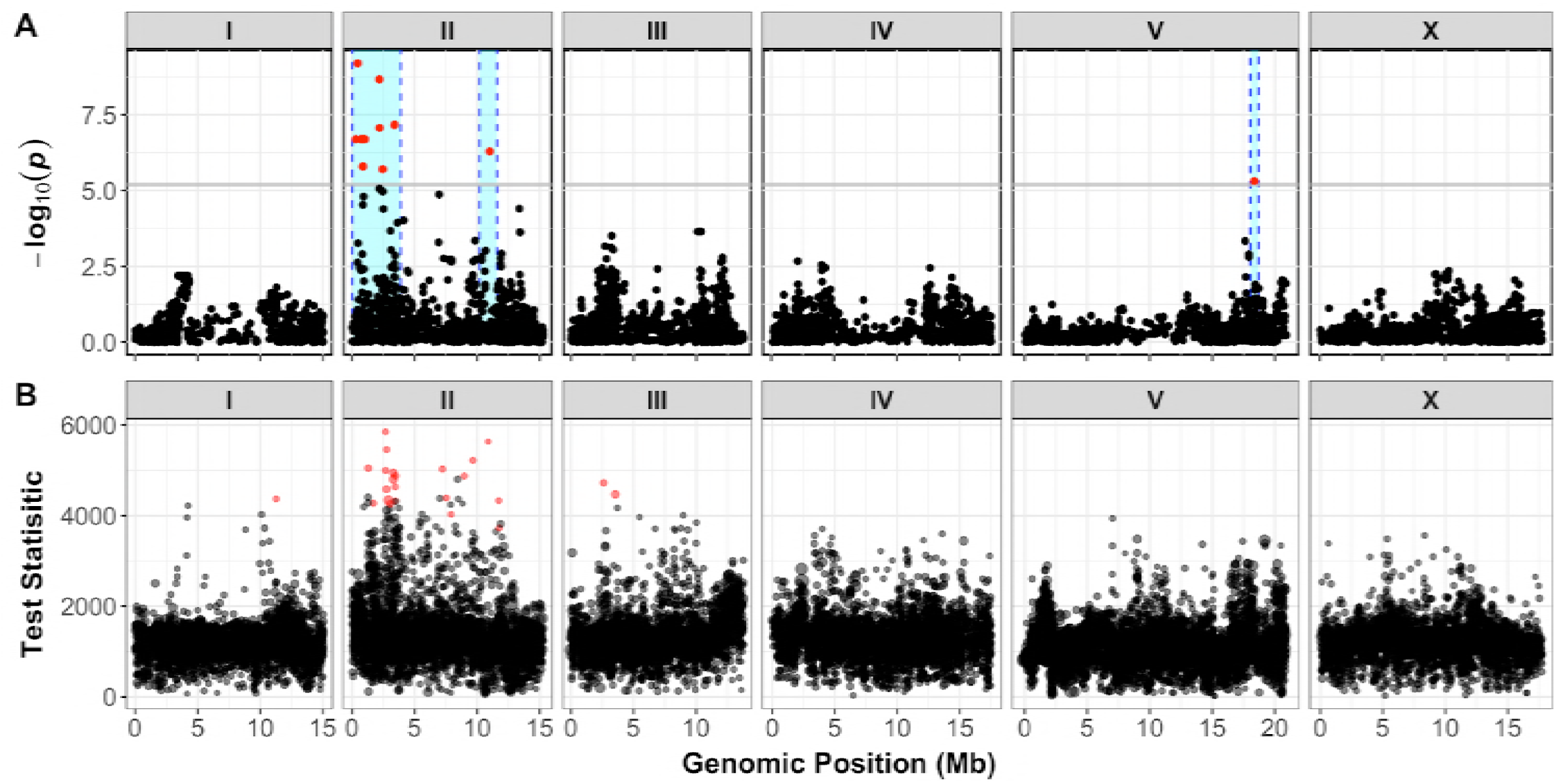
GWA mappings of 209 C elegans wild isolates in response to ABZ. A) The single-marker based Manhattan plot for animal length (q90.TOF) in the presence of ABZ is shown. Each dot represents an SNV that is present in at least 5% of the assayed population. The genomic location of each SNV is plotted on the x-axis, and the statistical significance of the correlation between genotype and phenotype is plotted on the y-axis. SNVs are colored red if they pass the genome-wide Bonferroni-corrected significance threshold, which is shown by a gray horizontal line. Genomic regions of interest are represented by cyan rectangles surrounding each QTL. In total, three QTL were identified, located on chr II and chr V. B) A genome-wide manhattan plot of animal length (q90.TOF) after ABZ exposure based on the gene-burden mapping approach is shown. Each dot represents a single *C. elegans* gene with its genomic location plotted on the x-axis, and the statistical significance of the correlation between genotype and phenotype plotted on the y-axis. Genes passing the burden test statistical significance threshold are colored in red. The burden test analysis identified genes significantly correlated with ABZ resistance on chr I, chr II, and chr III.

To complement the single-marker mapping described above, we performed a gene-burden approach to map BZ resistance [39,40]. Compared to the single-marker approach that includes SNVs with a minimum 5% minor allele frequency among all strains, the gene-burden approach incorporates all rare strain-specific variation within a gene for association testing [39,56]. With this alternative mapping strategy, we identified the same two QTL on chromosome II that we identified with the single-marker approach. In addition to the chromosome II QTL, we found a significant association between variation at the *ben-1* locus on chromosome III and animal-length variation in response to ABZ (Figure 1B). These results suggest that no variants are shared above 5% minor allele frequency among the wild isolates at the *ben-1* locus (chromosome III, 3,537,688-3,541,628 bp). An additional significant gene was detected on the right arm of chromosome I (Supplemental data 7 and 8).

### *C. elegans* ABZ resistance correlates with extreme allelic heterogeneity at the *ben-1* locus

To explain the differences between the single-marker and gene-burden based GWA mapping results with respect to *ben-1*, we investigated the natural variation found at this genomic locus in more detail (Figure 2; Supplemental data 9). We first used the *snpeff* function of the *cegwas* package [27,57] to look for SNVs in *ben-1* with predicted moderate-to-high effects on gene function [27,57]. These moderate-to-high impact variants include missense variants, splice donor and acceptor variants, and alternative start and stop codons that might disrupt the open reading frame of *ben-1*.

**Figure 2:**
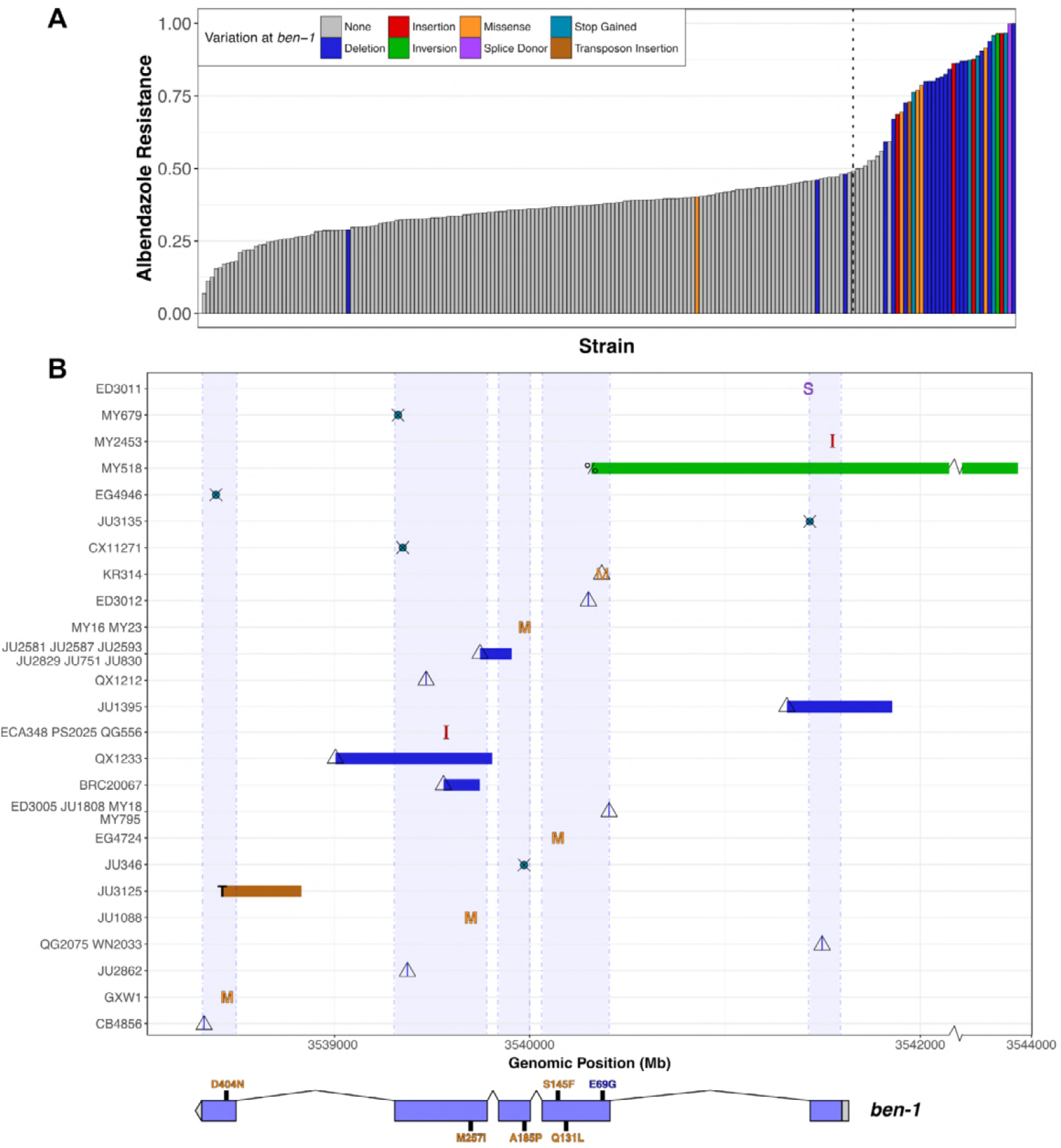
Variants in *ben-1* are highly correlated with ABZ resistance. A) A bar plot for the phenotypic response of *C. elegans* strains under ABZ exposure is shown. Each bar represents a single wild strain included in the HTA. Strains are sorted by their relative resistance to ABZ based on the mean animal length (q90.TOF) from four replicate measures. All strains that were found to have variants with predicted moderate-to-high impact and/or structural variation are colored by their specific type of variant. Strains similar to the N2 reference genome with respect to the *ben-1* locus are shown in grey. The black dotted line marks the point halfway point between the most and least resistant ABZ strains. B) An overview of variants found in the *ben-1* locus among *C. elegans* wild strains is shown. The genomic position of each variant is shown on the x-axis (S: splice donor variant; M: missense variant leading to amino acid substitution; crossed circle (⊗): alternative stop codon; triangle: deletion; I: insertion; percent sign: inversion; T: transposon insertion. Colors correspond to colors in (A). All strains with *ben-1* variants are listed on the y-axis in increasing order by their length in ABZ. For variants shared by two or more strains, the mean phenotype value of all corresponding strains determines their placement on the y-axis. A representation of the *ben-1* gene model is shown below the variant summary panel, including 5’ and 3’ UTRs (grey rectangle and bar), five exons (blue bars), and four introns (thin lines). Additionally, missense variants leading to amino acid substitutions are highlighted with their exact position and the corresponding amino acid exchange in the single letter code. Missense variants are colored orange if the corresponding strains do not contain any other *ben-1* variation. Missense variants that are always associated with deletion variants are colored blue.

In the set of 209 strains used for the GWA mapping, we identified 19 strains with moderate-to-high impact variants in the *ben-1* locus, as predicted by *snpEff* [57], including 13 strains with amino-acid substitutions. Amino-acid substitutions in β-tubulin genes are important markers for BZ resistance in parasitic nematodes and are hypothesized to reduce binding affinity of BZs to β-tubulin [16,17]. In the *C. elegans* panel of wild strains, we identified six strains that have the F200Y mutation in BEN-1, which is known to be the most common BZ resistance marker in livestock parasites [13]. Additionally, we detected novel amino-acid substitutions in BEN-1, which have not previously been associated with ABZ resistance in nematodes. These missense variants include an A185P substitution variant, found in two strains, as well as the E69G, Q131L, S145F, M257I, and D404N substitution variants, all of which were unique to single strains. In contrast to the amino-acid variants F200Y, E198A, and F167Y found in parasitic nematodes, which cluster at the putative ABZ binding site [58–60], these novel variants are more distributed throughout the protein structure (Supplemental Figure 3). We classified a strain as resistant to ABZ if its animal-length phenotype after ABZ exposure was greater than 50% of the difference between the most and least ABZ-resistant strains (Figure 2A). Eleven of the twelve strains with amino-acid substitutions are more resistant to ABZ than strains with no variation at the *ben-1* locus. Similarly, strains with predicted high-impact variants in *ben-1* are resistant to ABZ treatment. The strains with high-impact variants include five strains with unique premature stop codons and one strain with a predicted splice-donor variant at the end of exon 1. However, many ABZ-resistant strains in the *C. elegans* population do not contain any of these rare and common variants. Therefore, these strains either have other types of deleterious variants at the *ben-1* locus or are resistant to ABZ through a distinct mechanism. To differentiate these two possibilities, we manually curated all of the strains raw sequence read alignment files (BAM files), publicly available through the CeNDR website [27]. This in-depth investigation revealed extreme allelic heterogeneity at the *ben-1* locus in the natural *C. elegans* population. Twenty-seven strains had either rare deletions or insertions in *ben-1* (22 deletions, 5 insertions; Supplemental data 9). Interestingly, a subset of individuals with missense variants described above were found to be in perfect linkage disequilibrium (LD) with nearby deletion alleles. All six strains with the F200Y allele share the same deletion of approximately 160 bp that partially removes exons 3 and 4. The close proximity of the annotated F200Y to this 160 bp deletion suggested that the F200Y variant is actually an error in read alignment, which we confirmed by manual review of the BAM files of corresponding strains (JU751, JU830, JU2581, JU2587, JU2593, JU2829) (genome browser on CeNDR) [27]. The majority of paired-end reads at this genomic position that lead to the call of the F200Y allele in *ben-1*, map with their mate reads to the locus of the β-tubulin gene *tbb-2*, which is in relative close genomic distance (chromosome III, 4,015,769 - 4,017,643 bp). Additionally, the amino acid substitution (E69G), which is only observed in a single isotype (KR314), co-occurs with a deletion in exon 2 that is predicted to cause a frameshift in the *ben-1* open reading frame.

Of all the strains with *ben-1* SNVs and indels, only three individuals were not classified as resistant to ABZ treatment. The least resistant of these three strains was the CB4856 strain, which contains a nine base pair deletion near the end of the *ben-1* coding sequence. Because this in-frame deletion variant is at the end of the coding sequence, we hypothesized that CB4856 still contains a functional copy of the *ben-1* gene and this variant likely does not confer resistance to ABZ. The next least resistant strain is QG2075, which shares a four base pair deletion in exon 1 of *ben-1* with the ABZ-resistant isotype WN2033. During the growth phase of our HTA assay, we noted that the QG2075 strain has a slow-growth phenotype in normal growth conditions, which is likely confounding our phenotypic measurements in ABZ. Finally, the JU2862 strain has a unique four base pair deletion in exon 4 of *ben-1*, that is predicted to cause a frameshift in the *ben-1* open reading frame. We note that JU2862 is right at our arbitrary ABZ-resistance threshold and upon re-phenotyping with higher replication might be classified as ABZ resistant.

Among the remaining ABZ-resistant strains, we identified a putative transposon insertion in exon 5 of *ben-1* in the JU3125 strain. The genomic origin of this putative transposon insertion is from position 17.07 Mb on chromosome X and corresponds to a cut and paste DNA transposon Tc5B, which is part of the TcMar-Tc4 transposon superfamily [61,62]. Finally, the MY518 strain is resistant to ABZ but did not contain any of the above classes of variation. However, we did identify a 1 kb inversion that spans exon 1 and the promoter region of *ben-1*. Remarkably, all structural variants present in *ben-1* are only present in one or few wild strains and are mostly predicted to cause loss of *ben-1* function (Figure 2). Altogether, the putative loss-of-function variants described above explain 73.8% of the phenotypic variation present in the natural *C. elegans* population.

### Within-species selective pressures at the *ben-1* locus

The presence of multiple low-frequency *ben-1* alleles that confer resistance to ABZ treatment suggests that selection might have acted on this locus within the *C. elegans* population. To determine if this hypothesis is possible, we calculated the Ka/Ks ratio between *ben-1* genes from *C. elegans* and from two diverged nematode species *C. remanei* and *C. briggsae* [63]. The Ka/Ks ratio between the *ben-1* coding sequences of *C. elegans* and these two diverged species is **~**0.008, which indicates that the evolution of this locus is constrained across species. However, among the 249 wild isolates within the *C. elegans* species [27], we identified 10 synonymous and 22 nonsynonymous, stop-gained, or splice-site variants in the *ben-1* locus when compared to the N2 reference genome [25]. We note that the synonymous variants identified in *ben-1* are primarily found in wild strains that contain putative loss-of-function alleles. These results indicate that, despite the evolutionary constraint we observe at the *ben-1* locus across nematode species, an excess of potentially adaptive mutations (~2.2X nonsynonymous variants) have arisen within the *C. elegans* species. These results are consistent with our estimates of Tajima’s *D* [64] at the *ben-1* locus. When we considered all variant types across coding and non-coding regions of *ben-1*, we found Tajima’s *D* to be −2.02. When we only consider putative loss-of-function variants in the *ben-1* locus, the Tajima’s *D* estimate is −2.59. This large negative value of Tajima’s *D* likely reflects the high number of rare *ben-1* alleles present in the *C. elegans* population. We next calculated Tajima’s *D* for the genomic region surrounding *ben-1* (Figure 3A; Supplemental data 10). These results show a strong negative dip in Tajima’s *D* (< −1.5) between 3.50 and 3.55 Mb, which is the region directly surrounding *ben-1* (3.537-3.541 Mb). This observation indicates that the *C. elegans* population has undergone a population expansion or that the region surrounding *ben-1* may have been subject to selective pressures. To differentiate between these two possibilities, we calculated Fay and Wu’s *H* [65] and Zeng’s *E* [66] for *ben-1* (Figure 3A; Supplemental data 11). We noticed that these two statistics showed peaks around the *ben-1* locus, which do not correspond to our observations of multiple low minor allele frequency alleles. However, when we consider *ben-1* coding variation and non-coding variation separately, we found the *H_coding_* statistic to be 0.34 and the *H_noncoding_* statistic to be −5.5, which indicate that when considering high frequency derived alleles, the coding sequence of this locus is evolving neutrally and that there are many high-frequency derived alleles in the intronic and UTR regions. By contrast, we found the *E_coding_* statistic, which considers low and high frequency alleles, to be −1.7 and the *E_noncoding_* to be 3.4. Taken together, these results indicate that recent selective pressures have acted on the *ben-1* locus.

**Figure 3:**
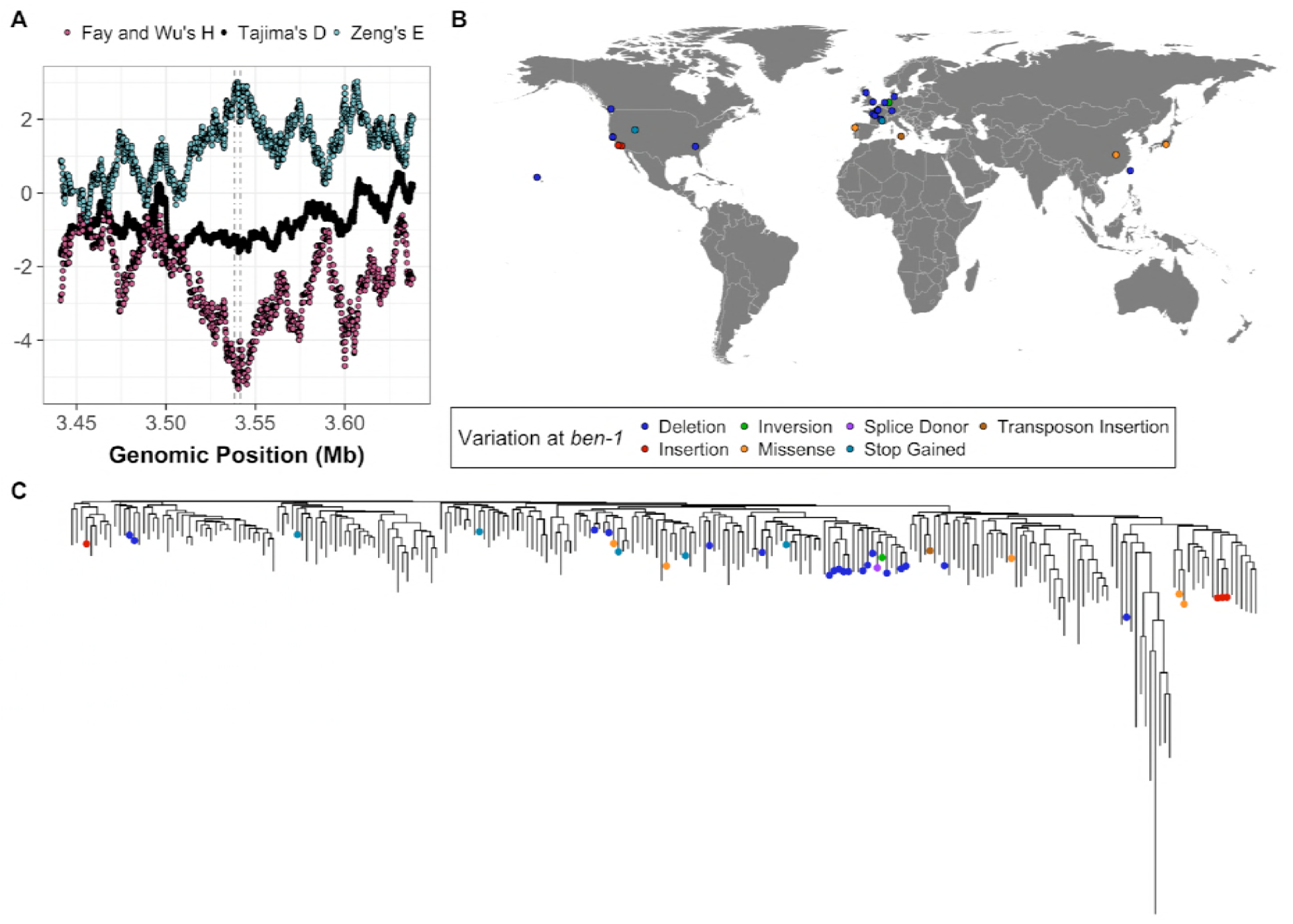
Population-level summary of ben-1 variants. A) Fay and Wu’s *H* (pink), Tajima’s *D* (black), and Zeng’s *E* (blue) for the genomic region surrounding *ben-1*. Genomic position is shown on the x-axis, and the value for each displayed statistic is shown on the y-axis. The neutrality statistics were calculated using a sliding window approach (100 SNVs window size and 1 SNV slide distance) using only SNV data. B) The global distribution of strains that contain moderate-to-high predicted variation in *ben-1*. Each dot corresponds to the sampling location of an individual strain and is colored by the type of variant discovered in the *ben-1* locus. C) The genome-wide phylogeny of 249 *C. elegans* strains showing that variation in the *ben-1* locus occurred independently multiple times during the evolutionary history of the species. The dots on individual branch nodes correspond to strains with variation in *ben-1* and have the same color code as in panel B.

If localized selective pressures, possibly caused by increased levels of environmental BZ, drove the loss of *ben-1* function in a subset of wild strains, then we could expect to see geographic clustering of resistant strains. When we looked at the distribution of strains with putative *ben-1* loss-of-function variants, we observed no trend in the sampling locations of these individuals (Figure 3B). However, we do note that highly diverse strains sampled from various locations on the Pacific Rim [27,37,38] do not harbor any putative *ben-1* loss-of-function alleles (Figure 3B). This diverse set of strains is hypothesized to represent the ancestral state of *C. elegans*, because they do not show signs of chromosome-scale selective sweeps throughout their genomes [38]. The lack of *ben-1* loss-of-function variants in these ancestral strains suggests that variation in *ben-1* arose after individuals in the species spread throughout the world. We next considered that strains with putative *ben-1* loss-of-function alleles might have been isolated from similar local environments and substrates. Strains that are predicted to have a functional *ben-1* gene have been isolated from a greater diversity of environmental sampling locations (Supplemental Figure 4A) and sampling substrates (Supplemental Figure 4B) than strains with predicted loss-of-function variants in *ben-1*. However, a hypergeometric test for enrichment within specific locations and substrates showed no signs of significant enrichment. This lack of enrichment might be caused by biases in global coverage of *C. elegans* sampling and low sampling density. Nevertheless, the observation that the diversity of *ben-1* alleles arose independently on various branches of the *C. elegans* phylogeny lends support to the hypothesis that local selective pressures have acted on the *ben-1* locus (Figure 3C).

### *ben-1* natural variants confer BZ resistance to sensitive *C. elegans* strains

The majority of the variants we observe at the *ben-1* locus are predicted to result in the loss of gene function. Our findings in *C. elegans* are in contrast to findings in parasitic nematode populations that describe the F200Y and other missense variants as the major variants contributing to BZ resistance [13]. To test whether the putative loss-of-function variants in *ben-1* that we observe in the *C. elegans* population confer the same level of BZ resistance as the known F200Y allele, we introduced a *ben-1* deletion in the N2 strain using CRISPR/Cas9 (referred to as Del). In addition to the Del strain, we introduced the F200Y *ben-1* allele into the N2 strain (referred to as F200Y) to test whether this allele confers BZ resistance in *C. elegans* and to compare the levels of BZ resistance between the loss-of-function and missense alleles. We exposed the N2 parental, Del, and F200Y strains to 12.5 μM ABZ using the HTA described above. Both the Del and F200Y variants conferred ABZ resistance to the otherwise sensitive N2 parental strain (Figure 4A; Supplemental Figure 5; Supplemental Data 12 and 13). Interestingly, we found no significant difference in BZ resistance between the Del and F200Y strains (*p*-value = 0.99, TukeyHSD). These results show that putative loss-of-function variants of *ben-1* and known parasitic BZ resistance alleles confer BZ resistance in otherwise isogenic *C. elegans* genetic backgrounds.

**Figure 4:**
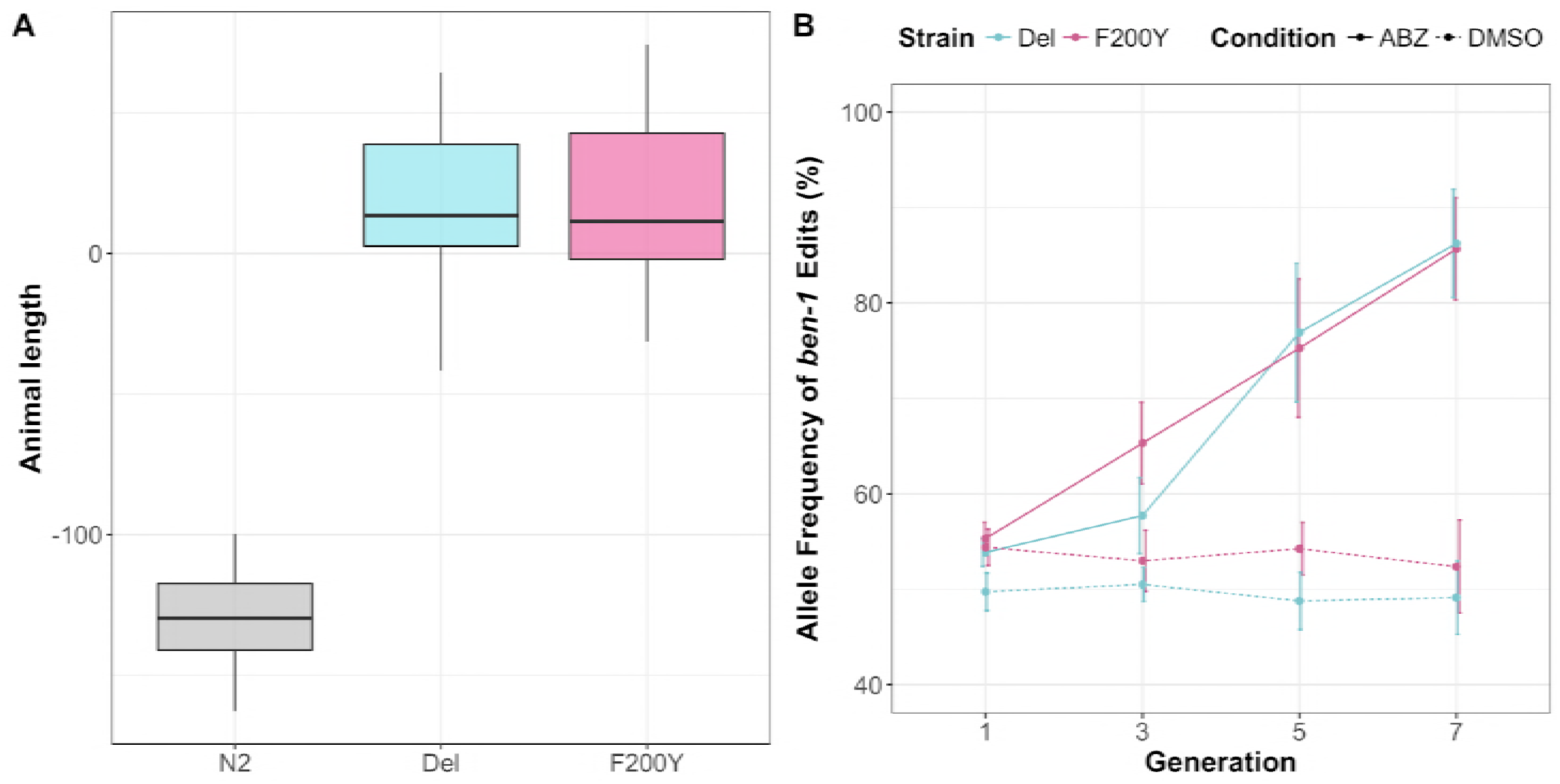
F200Y allele replacement and *ben-1* deletion confer resistance to the ABZ sensitive N2 strain. **A)** Tukey box plots of the animal length phenotypes for at least 100 replicates of the generated *ben-1* allele-replacement strains after ABZ treatment is shown. The y-axis represents the animal length phenotype after correcting for growth in DMSO conditions. Both *ben-1* allele strains are significantly more resistant to ABZ treatment than the N2 strain (*p*-value < 1E-10, TukeyHSD). **B)** The results from a multi-generation competition experiment between a barcoded N2 strain and the *ben-1* allele-replacement strains are shown. The generation number is shown on the x-axis, and the allele frequency of the *ben-1* allele-replacement strains is shown on the y-axis.

To complement the results from the liquid-based HTA experiments, we performed a plate-based competition assay. In the competition experiments, we individually competed the F200Y, Del, and parental N2 strains against an N2 strain that contains a barcode sequence (PTM229). We quantified the relative allele frequencies of the barcoded strain for the first, third, fifth, and seventh generations of the competition assay (Figure 4B; Supplemental Data 14 and 15). Throughout the competition assay, and for all strains tested, the relative frequencies of the barcoded strain did not significantly deviate from the initial frequency when grown on DMSO plates. These results suggest that in standard laboratory conditions, the BEN-1 F200Y and *ben-1* deletion alleles do not have fitness consequences. We observed the same trend when we competed the N2 and the barcoded N2 strains on ABZ plates. However, the allele frequencies of the barcoded strain dropped to ~20% when competed against strains that contain either of the two *ben-1* alleles on ABZ plates (relative fitness w = ~ 1.3, both). These two independent assays show that the two *ben-1* alleles confer BZ resistance with no negative fitness consequence under standard laboratory growth conditions.

### Additional genomic intervals contribute to ABZ resistance in the *C. elegans* population

Above, we showed that extreme allelic heterogeneity at the *ben-1* locus explains 73.8% of the phenotypic variation in response to ABZ treatment. To identify potential genomic loci that contribute to the remaining 26.2% of the observed BZ response variation, we statistically corrected the animal length phenotype in response to ABZ treatment by using the presence of a putative loss-of-function variant at the *ben-1* locus in a strain as a covariate for linear regression. After correcting for variation at the *ben-1* locus, we performed GWA mapping using both the single-marker (Figure 5A; Supplemental Data 16, 17, and 18) and gene-burden (Supplemental Figure 6; Supplemental Data 19, 20, and 21) approaches. Both of these approaches resulted in the disappearance of the two QTL on chromosome II and the QTL on chromosome V (Figure 5A, Supplemental Figure 7). This result suggests that the cumulative variation at the *ben-1* locus is in complex interchromosomal linkage disequilibrium (LD) with these three loci (Supplemental Figure 7), despite the loci on chromosomes II and V not being in strong LD with each other (Supplemental Figure 2B). Interestingly, using the single-marker GWA-mapping approach, we found an additional genomic locus on chromosome X that was significantly associated with ABZ resistance (Figure 5A; Supplemental Figure 8). However, the gene-burden based approach did not result in significant associations between ABZ resistance and rare variation within any genes. These results suggest that ABZ resistance in the *C. elegans* population is caused by at least two loci because additional genetic variation present in the *C. elegans* population contributes to phenotypic variation in response to ABZ treatment.

**Figure 5:**
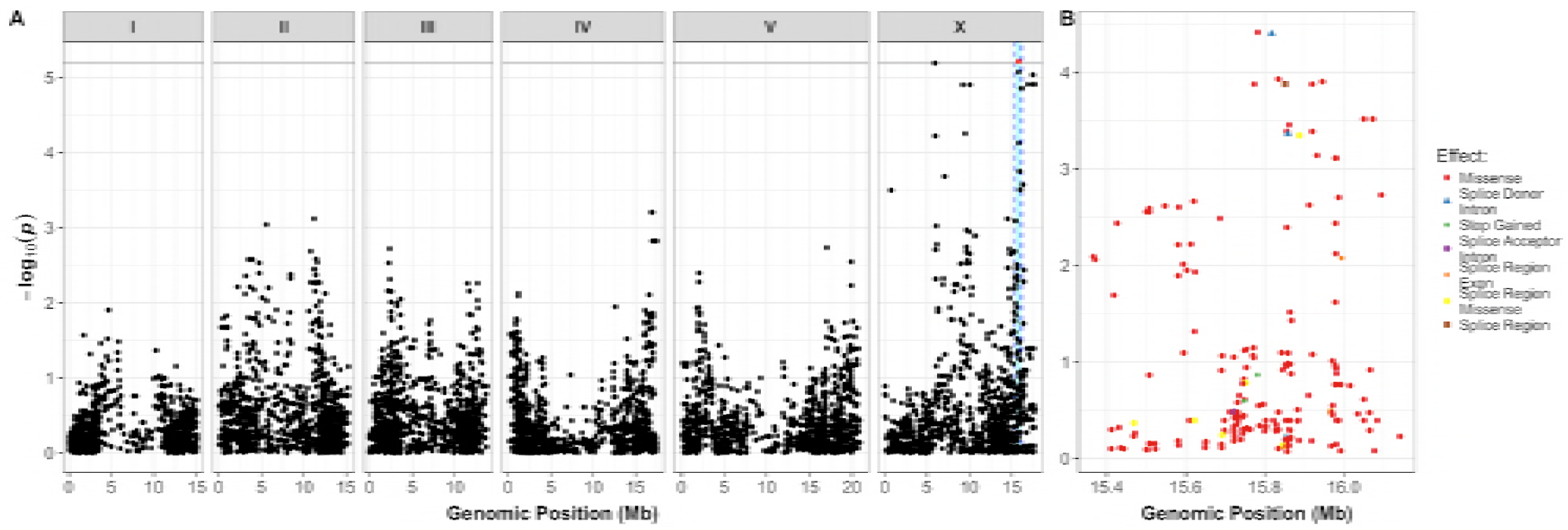
Regression of the putative *ben-1* LoF variants identifies a novel QTL on chromosome X. A) Manhattan plot of the single-marker based GWA mapping showing that a nominally significant QTL on chromosome X is associated with residual phenotypic variation in response to ABZ treatment. Each dot represents an SNV that is present in at least 5% of the assayed population. The genomic location of each SNV is plotted on the x-axis, and the statistical significance is plotted on the y-axis. SNVs are colored red if they pass the Bonferroni-corrected significance threshold, which is shown by a gray horizontal line. The genomic region of interest is represented by cyan rectangle surrounding the chromosome X QTL. B) Fine-mapping of the chromosome X QTL showing all variants with moderate-to-high predicted effects within the chromosomes X QTL region of interest. The genomic position in Mb is shown on the x-axis, and the –*log*10(*p*) value, which represents the strength of the association between residual ABZ resistance and the plotted variant, is shown on the y-axis. Dots are colored by the type of variant.

## Discussion

ABZ is a broadly administered BZ used to treat parasitic nematode infections in humans and livestock [4,67]. Already a significant problem in veterinary medicine, the heavy reliance of ABZ and other BZ compounds in MDA programs, the small repertoire of other anthelmintic compounds, and high rates of re-infection of parasitic nematodes in endemic regions around the world have raised the fear of the emergence of BZ resistance among parasitic nematode populations that infect humans [4,68]. Though the BZ-resistance alleles found in model organisms through classical genetic screens have had little direct translatability to parasitic nematodes, these classic alleles have been instrumental toward the elucidation of the mechanism of action of BZs [4,69]. Recent advances in sequencing technologies have enabled researchers to take a quantitative genetics approach to search for novel mechanisms of anthelmintic resistance in natural parasitic and non-parasitic nematodes [27,29,30,70–72]. In the present study, we leveraged genetic diversity within the *C. elegans* population to study the genetic basis of ABZ resistance within this species.

To identify the genetic basis of ABZ resistance in *C. elegans*, we used two unbiased genome-wide association approaches, single-marker [35,36] and gene-burden [40] based mappings. We found that the *C. elegans* population harbors extreme allelic heterogeneity at the *ben-1* locus, and that this variation contributes to differential ABZ resistance among wild isolates (Figure 1B, Figure 2A). The variants in the *ben-1* locus that contribute to ABZ resistance are present at low allele frequencies (<0.05 MAF) within the global *C. elegans* population and arose independently during the evolutionary history of the species (Figure 3). As a result of this complex demographic history, common variants (>0.05 MAF) used as genetic markers in the mappings near the *ben-1* locus are not in LD with the *ben-1* alleles that confer ABZ resistance. Therefore, we were unable to detect an association between the *ben-1* locus and ABZ response with the single-marker based mapping approach. We note that burden-based mapping approaches are only possible in species with well annotated genomes and genome-wide genetic variation, such as *C. elegans* and other model organisms. To extend this type of analysis to parasitic species, more effort will need to be made toward improving parasitic nematode reference genomes and the accumulation of genome-wide variation resources [73,74]. In addition to the strong association between ABZ responses and variants in the *ben-1* locus, we identified a novel ABZ-response QTL on chromosome X after accounting for variants in the *ben-1* locus (Figure 5). The identification of this QTL suggests that mechanisms independent of β-tubulin could account for BZ resistance in the *C. elegans* and in parasitic nematode populations.

The majority (22/25) of *ben-1* alleles that correlate with ABZ resistance are predicted to result in the loss of *ben-1* function (Figure 2). These results suggest that the function of the *ben-1* gene is not essential to the survival of *C. elegans* in natural habitats. This observation is in close agreement with previous work using the laboratory-derived N2 strain, which showed that putative *ben-1* loss-of-function alleles do not confer observable defects in standard laboratory growth conditions [12]. However, our analysis of Ka/Ks between the *C. elegans ben-1* and the distantly related *C. briggsae* and *C. remanei* β-tubulin coding sequences suggest that evolution of this gene is highly constrained. A possible explanation for this discrepancy is that the *ben-1* gene has a highly specialized function in natural settings and that survival in the presence of a strong selective pressure, such as the presence of increased concentrations of environmental BZs, outweighs the necessity of this specialization. A possible source of environmental BZs are microbes that *C. elegans* might contact in their natural habitat. For example, 5,6-dimethylbenzimidazole is a BZ derivative that is produced naturally by prokaryotes [75]. However, whether these BZ derivatives bind to and inhibit β-tubulins remains unclear. Alternatively, natural selection on *C. elegans ben-1* might occur by contamination of nematode niches by synthetic BZs produced by humans. Initially developed as fungicides in the 1960s, several BZs, including thiabendazole, benomyl, and carbendazim are extensively used to treat crops, fruits, and grains [76]. A third source of environmental BZs might be runoff from livestock farms because they are used extensively to prevent parasite infections in these animals [77–80]. The observation that putative *ben-1* loss-of-function alleles are only found at low frequencies in the *C. elegans* population might suggest that individuals with these alleles are eventually removed from the population, though we see no evidence for a decrease in fitness in laboratory competitions experiments, which might not recapitulate natural settings (Figure 4). Taken together, our observations indicate that the independent putative *ben-1* loss-of-function alleles might have arose recently within the *C. elegans* population. Considering these findings in retrospect, it is extremely fortunate that N2 was used for initial studies of BZ sensitivity. Rapid progress on the mechanism of action for BZ compounds from the free-living *C. elegans* model to parasites might have not occurred otherwise, strongly illustrating the pitfalls associated with the study of a single genetic background to elucidate anthelmintic resistance.

The remaining (3/25) *ben-1* alleles we found to be correlated with ABZ resistance in the *C. elegans* population result in missense variants. These missense variants are S145F, A185P, and M257I and are found in one, two, and one *C. elegans* strain, respectively. Remarkably, substitutions of the amino acid residues A185 (A185S) and M257 (M257L) were previously described to confer BZ resistances in *Tapesia yallundae* and *Aspergillus nidulans*, respectively [81,82]. Of these two previously identified residues, M257 is postulated to directly interact with BZs, because it is in close three-dimensional proximity to the known BZ interacting F200 residue [60]. The S145F allele is in close proximity to the highly conserved GGGTGS motif of the GTP binding and hydrolysis site [83]. A fourth missense mutation Q131L, which is present in a strain that was not phenotyped in our mapping study because it grows slowly under normal laboratory conditions, is located near the β-tubulin/α-tubulin interaction interface [84,85]. Upon re-phenotyping this isotype with the Q131L variant, we found that it is resistant to ABZ treatment (Supplemental Figure 9). A final missense variant (D404N) present in the *C. elegans* population was not correlated with ABZ resistance (Figure 2). Though theoretical structure-based evidence suggests that most of the missense variants in the *C. elegans* population associated with ABZ resistance cause a non-functional β-tubulin, further experiments are necessary to confirm these results.

The high prevalence of putative *ben-1* loss-of-function variants in the natural *C. elegans* population stands in stark contrast to the relative paucity of allelic diversity found in orthologous β-tubulin genes in parasitic nematodes. Perhaps parasitic nematode species do not have similarly high levels of functional redundancy of β-tubulins as has been observed in *C. elegans*. The main *C. elegans* β-tubulins are *tbb-1* and *tbb-2*, which are both ubiquitously highly expressed, functionally redundant in laboratory conditions, and contain a tyrosine at position 200 [86–88]. The co-expression of *tbb-1*, *tbb-2*, and *ben-1* in the nervous system [89] suggests that only one β-tubulin with F200 is required to confer sensitivity to BZs. Two other genes that encode for β-tubulins in *C. elegans* are *mec-7* and *tbb-4*. Both genes are only expressed in a subset of neuronal cells involved in chemo- and mechanosensation, and both have a phenylalanine at amino acid position 200 [88, 90–92]. *tbb-6* encodes for a highly diverged *C. elegans* β-tubulin (Supplemental Figure 10) that could be a stress-resistant tubulin because it is highly expressed during the unfolded protein response [93]. Despite having phenylalanine at amino acid position 200, these tubulins (TBB-4, MEC-7, and TBB-6) are likely not involved in BZ response because we see no difference in ABZ-resistance between control and ABZ conditions for BZ-resistant strains (Supplemental Data 12). Of the six β-tubulin genes described above, we only observe variation in *ben-1* and *tbb-6*, both of which have low-frequency variants with high predicted functional effects. These observations are also shown by estimates of Tajima’s *D* at these loci (Supplemental Figure 11).

The β-tubulin repertoire among parasitic nematode species is highly diverse likely because of multiple gene duplication events [94]. For example, *H. contortus* has four known β-tubulin genes, of which *Hco-tbb-iso-1* and *Hco-tbb-iso-2* are thought to be the targets of BZ, and *Hco-tbb-iso-3* and *Hco-tbb-iso-4* are closely related in sequence and expression pattern to *C. elegans tbb-4* and *mec-7* [94]. Despite the presence of a phenylalanine at amino acid position 200 in all four of the β-tubulins, the focus of the parasitology community is on missense variants present in *Hco-tbb-iso-1*. However, our data argue that it is important to consider the high level of sequence similarity among the β-tubulin genes. We see that nearby structural variants, like deletions, can alter sequence read alignments and misannotate orthologous β-tubulin gene sequences to artifactually create these variant sites. Careful read alignments and high read depth will be necessary to interpret variation across highly divergent parasites with orthologous β-tubulin genes. Additionally, similar to our observations of *ben-1* in *C. elegans*, deletion alleles of *H. contortus Hco-tbb-iso-2* have been observed in few field studies, but their importance for BZ resistance remains unclear [95,96]. This observation led Kwa and colleagues [95,97] to hypothesize that BZ resistance requires two steps. First, mutation of F200 to tyrosine in one β-tubulin isoform, and second, deletion of the second β-tubulin that contains F200. This hypothesis aligns well with our observation that the single F200Y change in *C. elegans* BEN-1 confers BZ resistance in the presence of TBB-1 (Y200) and TBB-2 (Y200). An additional layer of complexity comes from recent evidence in *Trichuris trichiura* that non-BEN-1 β-tubulins might also play a role in BZ resistance in diverse parasitic species [98]. Our results showing that BZ resistance is a complex trait within the *C. elegans* population (Figure 5) and others’ similar observations within parasitic nematode populations necessitates further investigation into the genetic and molecular mechanisms that underlie this trait.

## Material and Methods

### *C. elegans* strains

Animals were cultured at 20°C on modified nematode growth medium (NGM), containing 1% agar and 0.7% agarose [31] and seeded with OP50 *E. coli*. Prior to each assay, strains were passaged for at least four generations without entering starvation or encountering dauer-inducing conditions [31]. For the genome-wide association (GWA) studies, 249 wild isolates from CeNDR (version 20170531) were used as described previously [27,32]. Construction of *ben-1* allele-replacement and *ben-1* deletion strains are described in the corresponding section. All strain information can be found in Supplemental table 1.

### High-throughput fitness assay

The high-throughput fitness assays (HTA) were performed as described previously [32] with the exception that each strain was assayed in four technical replicates that consisted of four independent bleach synchronization steps across two days with two independent drug and control preparations. In short, strains were propagated for four generations on agar plates, followed by bleach synchronization. The embryos were titered to 96-well microtiter plates at a final concentration of approximately one embryo per microliter of K medium [33] with modified salt concentrations (10.2 mM NaCl, 32 mM KCl, 3 mM CaCl_2_, 3 mM MgSO_4_). After overnight incubation, hatched L1 larvae were fed with 5 mg/mL HB101 bacterial lysate (Pennsylvania State University Shared Fermentation Facility, State College, PA) and cultured for two days until they reached the L4 larval stage. Using the large particle flow cytometer COPAS BIOSORT (Union Biometrica, Holliston MA), three L4 larvae per well were sorted into new microtiter plates containing modified K medium, 10 mg/mL HB101 lysate, 50 μM kanamycin, and either 12.5 μM albendazole (ABZ) dissolved in 1% DMSO or 1% DMSO alone as a control. During the following four-day incubation, animals were allowed to mature and to produce offspring. Fitness parameters, including traits for animal length and brood size, were measured for each population under drug and control conditions using the COPAS BIOSORT platform after 96 hours. To facilitate body straightening for more accurate length determinations, animals were treated with sodium azide (50 mM) immediately before measurement.

### Processing of fitness traits for genetic mapping

Fitness parameters measured by the COPAS BIOSORT platform were processed using the R package *easysorter* [34] with some modifications to incorporate technical replicates into the analysis. The *easysorter* package is specifically developed for this type of data and includes reading and modifying of the raw data, pruning of anomalous data points, and regression of control and experimental phenotypes. Measured parameters include time-of-flight (animal length), extinction (optical density), and total object count (brood size) for each well. In short, the function *read_data* reads in raw phenotype data and runs a support vector machine to identify and eliminate air bubbles, which can be confused with nematodes. In the next step, the function *remove_contamination* removes data obtained from microtiter wells that were manually identified to contain bacterial or fungal contamination. To generate summarized statistics for each well, the function *sum_plate* calculates the 10^th^, 25^th^, 50^th^, 75^th^, and 90^th^ quantiles for all fitness parameters obtained. In this process, brood size is normalized by the number of animals sorted originally in each well. Next, biologically impossible data points passing certain cut-offs (n > 1000, n < 5, norm.n > 350) are eliminated (function *bio_prune*). We removed outlier replicates if they were outside 1.8 times the standard deviation of that strain’s median phenotype of a particular trait. To adjust differences among replicates of each assay, a linear model was applied using the formula (phenotype ~ experiment + assay), which replaced the *easysorter* function *regress* (*assay = TRUE*). The experiment component of the linear model corresponds to two independent drug preparations of the same strains and the assay component corresponds to blocks of different strains performed across multiple weeks. In addition, the *bamf_prune* function of the *easysorter* package was skipped, because previous outlier removal based on technical replicates made this step unnecessary. Finally, drug-specific phenotype data is calculated using the *regress* (*assay = FALSE*) function of the *easysorter* package. This function fits a linear model with the formula (phenotype ~ control phenotype) to account for any differences in population parameters present in the control DMSO-only conditions.

### Dose-response experiments

Albendazole (Sigma Aldrich Product #A4673) (ABZ) dose-response experiments were performed on a set of four genetically divergent *C. elegans* strains, including the laboratory strain N2 and three different wild isolates (CB4856, JU775, DL238), to determine suitable drug concentrations for subsequent GWA experiments. To this end, strain-specific drug responses were measured in four technical replicates using the HTA as described above at concentrations of 3.125 μM, 6.25 μM, 12.5 μM, and 25 μM ABZ. A suitable drug concentration for subsequent GWA experiments was selected based on the lowest concentration in which we observed a significant difference in ABZ response among strains for brood size and animal length (Supplemental data 1 and 2).

### Genome-wide association (GWA) mappings

The GWA mappings were performed on the processed ABZ HTA phenotype data of 209 *C. elegans* wild isolates. In short, the *easysorter* processed phenotype data was analysed using the *cegwas* R package for association mapping [27]. This package uses the EMMA algorithm for performing association mapping and correcting for population structure [35], which is implemented by the *GWAS* function in the rrBLUP package [36]. In detail, the GWAS function was used with the following command: rrBLUP::GWAS (min.MAF = 0.05, P3D = FALSE). The kinship matrix used for association mapping was generated using whole-genome high-quality single-nucleotide variants (SNVs) [37] and the *A.mat* function from the rrBLUP package. All SNVs included in the marker set for GWA mapping had a minimum 5% minor allele frequency in the 240 isotype set [38]. Quantitative trait loci (QTL) were defined by at least one SNV that passed the Bonferroni-corrected threshold and were processed further using fine mapping, as described previously [32]. Computational fine mapping of the genomic regions of interest was performed as described previously [32]. We used the following linear model to correct for the presence of a putative *ben-1* LoF variant: *lm*(*animal length*~(*ben* – 1 *LoF*)). The list of strains that were considered to have putative LoF variants and the *ben-1-*corrected phenotype data are presented (Supplemental data 16). We did not include CB4856 as a strain with a putative LoF variant because it has a 9 bp in-frame deletion near the end of the gene. We performed single-marker mappings as described above and gene-burden mappings as described below using these regressed phenotypes (Supplemental data 17-21).

### Burden Testing

Burden test analyses were performed using RVtests [39] and the variable-threshold method [40]. We called SNV using bcftools [41] with settings previously described [27,37,39]. We next performed imputation using BEAGLE v4.1 [42] with *window* set to 8000, *overlap* set to 3000, and *ne* set to 17500. Within RVtests, we set the minor allele frequency range from 0.003 to 0.05 for burden testing.

### Generation of *ben-1* allele replacement and deletion strains

All *ben-1* edited strains assayed in this study were generated in the N2 background by CRISPR/Cas9-mediated genome editing, using a co-CRISPR approach [43] and Cas9 ribonucleoprotein (RNP) delivery [44]. For the *ben-1* F200Y allele replacement strains, sgRNAs for *ben-1* and *dpy-10* were ordered from Synthego (Redwood City, CA) and injected at final concentrations of 5 μM and 1 μM, respectively. Single-stranded oligodeoxynucleotides (ssODN) templates for homology-directed repair (HDR) of *ben-1* and *dpy-10* (IDT, Skokie IL) were used at final concentrations of 6 μM and 0.5 μM, respectively. Cas9 protein (IDT, Product #1074182) was added to the injection mixture at a concentration of 5 μM and incubated with all other components for ten minutes at room temperature prior to injection. All concentrations used for the sgRNA-mediated allele replacement were adapted from the work of Prior and colleagues [45]. N2 *ben-1* deletion strains were generated using two *ben-1* specific crRNAs synthesized by IDT (Skokie, IL), which targeted exon 2 and exon 4. For the injection mixture, *ben-1* crRNAs were used at final concentration of 8.3 μM each, mixed with *dpy-10* crRNA and tracrRNA (IDT, Product #1072532) at final concentrations of 1.2 μM and 17.6 μM, respectively, and incubated at 95°C for five minutes. After cooling to room temperature, Cas9 protein was added at a final concentration of 15.25 μM (IDT Product #1074181) and incubated for five minutes before *dpy-10* ssODN was added to a concentration of 5 μM.

RNP injection mixtures were microinjected into the germline of young adult hermaphrodites (P0) and injected animals were singled to fresh 6 cm NGM plates 18 hours after injection. Two days later, F1 progeny were screened, and animals expressing a Rol phenotype were transferred to new plates and allowed to generate progeny (F2). Afterwards, F1 animals were genotyped by PCR. For the *ben-1*(F200Y) allele replacement, ssODN repair templates contained the desired edit and a conservative change of the PAM site to prevent sgRNA:Cas9 cleavage. In addition, the PAM change introduced a new restriction site. PCRs were performed using the primers oECA1297 and oECA1298, and PCR products were incubated with BTsCI restriction enzyme (R0647S, New England Biolabs, Ipswich, MA) to identify successfully edited animals by differential band patterns. For genotyping of *ben-1* deletion strains, the primers oECA1301 and oECA1302 were used and successful deletions were identified by shorter PCR products. Non-Rol progeny (F2) of F1 animals positive for the desired edits were propagated on separate plates to generate homozygous progeny. F2 animals were genotyped afterwards, and PCR products were Sanger sequenced for verification. Generated *ben-1*(F200Y) replacement strains and *ben-1* deletion strains were phenotyped for ABZ response using the HTA as described above. All oligonucleotide sequences are listed in the supplement (Supplemental table 2).

### Competition assays

Pairwise multi-generation competition assays were performed between a *ben-1* wild-type strain PTM229 and the *ben-1* edited strains, containing either the F200Y allele replacement or a *ben-1* deletion. All strains were generated in the N2 background. For the assay, strains were bleach-synchronized and embryos were transferred to 10 cm NGM plates. 48 hours later, seven L4 larvae per strain were transferred to 50 fresh 6 cm NGM plates containing either 1.25 μM ABZ for drug selection or DMSO as control. The ABZ concentration was determined by dose-response assays beforehand as the lowest concentration that caused a developmental delay in PTM229 and N2 when added to NGM. For each strain combination, 50 plates were grown, representing 10 technical replicates of five independent populations. Plates were grown for one week until starvation. Animals were transferred to fresh plates by cutting out a 0.5 cm^3^ agar chunk. After every culture transfer, starved animals were washed off the plates with M9, and DNA was collected using the Qiagen DNeasy Kit (Catalog #69506). Allele frequencies of PTM229 compared to wild-type N2 or the *ben-1* edit strains in each replicate populations were measured using Taqman analysis in a Bio-Rad QX200 digital droplet PCR system. Digital PCR was performed following the standard protocol provided by Bio-Rad with the absolute quantification method. To calculate the relative fitness w of the competitive strains, we used linear regression to fit the relative allele frequencies at the first, third, fifth, and seventh weeks into a one-locus genic selection model [46]. All oligonucleotide sequences are listed in the supplement (Supplemental table 2).

### Computational modelling of *ben-1* variants

We obtained the predicted BEN-1 peptide sequence from WormBase [25]. We used the online utility PHYRE2 to generate a homology model of the predicted BEN-1 peptide [47]. We specified the intensive homology search option for running PHYRE2. Visualization of the BEN-1 homology model and highlighting of the variants of interest was performed using PyMol [48].

### Statistical analyses

Phenotype data are shown as Tukey box plots and p-values were used to assess significant differences of strain phenotypes in allele-replacement experiments. All calculations were performed in R using the *TukeyHSD* function on an ANOVA model with the formula (*phenotype ~ strain*). P-values less than 0.05 after Bonferroni correction for multiple testing were considered to be significant.

### Population Genetics

Sliding window analysis of population genetic statistics was performed using the PopGenome package in R [49,50]. All sliding window analyses were performed using the imputed SNV VCF available on the CeNDR website with the most diverged isotype XZ1516 set to the outgroup [27,42,51]. Window size was set to 100 SNVs with a slide distance of one SNV. The Tajima’s *D* calculation, using all of the manually curated variants in the *ben-1* locus, was performed using modified code from the developers of the LDhat package [52]. Estimates of Ka/Ks were performed using an online service (http://services.cbu.uib.no/tools/kaks). Multiple sequence alignments were performed using MUSCLE [53] through the online service (https://www.ebi.ac.uk/Tools/msa/muscle/). The β-tubulin neighbor-joining phylogeny construction was performed using Jalview 2 [54]. Linkage disequilibrium (LD) of QTL markers was calculated using the *genetics* [55] package in R. LD calculations are reported as *r* = – *D / sqrt*(*p*(*A*) * *p*(*a*) * *p*(*B*) * *p*(*b*))where *D* = *p*(*AB*) – *p*(*A*) * *p*(*B*).*r* = −*D* / *sqrt(p(A) * p(a) * p(B) * p(b))*

## Acknowledgements

We thank Shannon C. Brady, Kyle Siegel, and members of the Andersen lab for editing the manuscript for flow and content. We thank members of the Andersen lab for making reagents used in the experiments presented in the manuscript.

## Supplemental Data

**TS1**: HTA ABZ dose response data_raw(.Rda)

**TS2**: HTA ABZ dose response data_processed(.tsv)

**TS3**: HTA phenotype data of *C. elegans* wild strains_raw(.Rda)

**TS4**: HTA phenotype data of *C. elegans* wild strains_processed(.tsv)

**TS5**: GWA single marker mapping data_processed(.tsv)

**TS6**: GWA single marker fine mapping data(.tsv)

**TS7:** GWA gene-burden mapping phenotype data(.ped)

**TS8**: GWA gene-burden mapping data(.tsv)

**TS9**: Positions and effects for all *ben-1* variants(.tsv)

**TS10**: Summary of population genetics data(.Rda)

**TS11**: Tajima’s D, Fay’s Wu and Zeng’s E for *ben-1* (.tsv)

**TS12:** HTA phenotype data of *C. elegans ben-1* mutants_summarized(.tsv)

**TS 13**: HTA phenotype data of *C. elegans ben-1* mutants_processed(.tsv)

**TS 14**: competition assay data_raw(.tsv)

**TS 15**: relative fitness calculations(.xml)

**TS 16**: *ben-1* regressed with *ben-1* covariate(.tsv)

**TS 17**: GWA single-marker mapping data after *ben-1* regression(.tsv)

**TS 18**: GWA single-marker fine mapping data after *ben-1* regression(.tsv)

**TS 19:** GWA gene-burden mapping phenotype data after ben-1 regression(.ped)

**TS 20:** GWA gene-burden mapping data after ben-regression(.assoc)

**TS 21:** GWA gene-burden mapping data after ben-1 regression(.tsv)

**TS 22**: Tajima’s D for other *C. elegans* beta-tubulins (.tsv)

**TS 23**: HTA re-phenotyping of *C. elegans* wild strains(.Rda)

**Supplemental Table 1**: strains(.csv)

**Supplemental Table 2**: Oligonucleotide sequences(.csv)

**Supplemental Figure 1:**
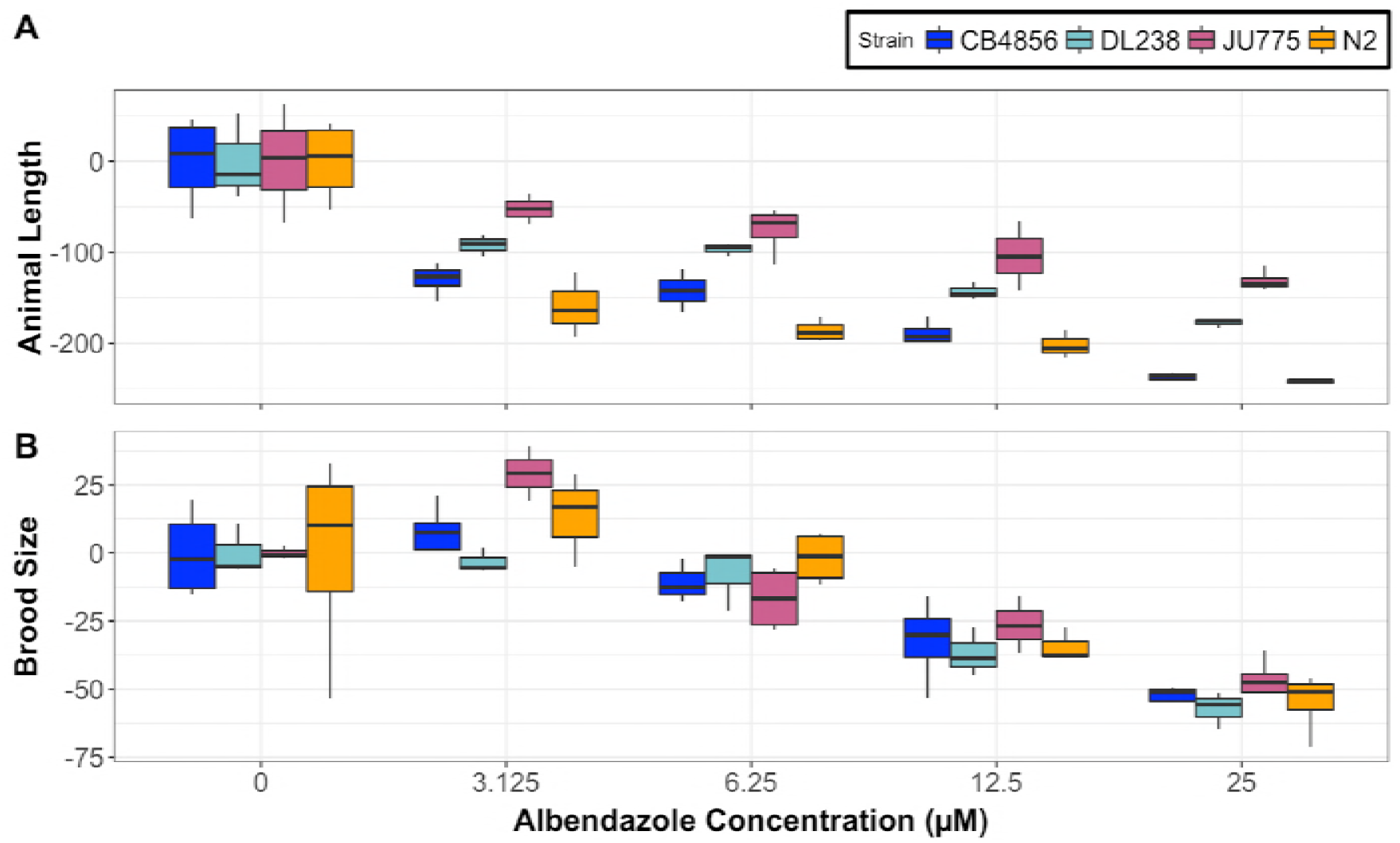
Dose response of four genetically divergent *C. elegans* strains after ABZ exposure. The box plots show a representative dose-response experiment on four *C. elegans* strains for A) animal length (q90.TOF) and B) brood size (norm.n). The ABZ concentration is plotted on the x-axis, and individual replicate trait values subtracted from the mean in DMSO control conditions is plotted on the y-axis. Each box represents four technical replicates.

**Supplemental Figure 2:**
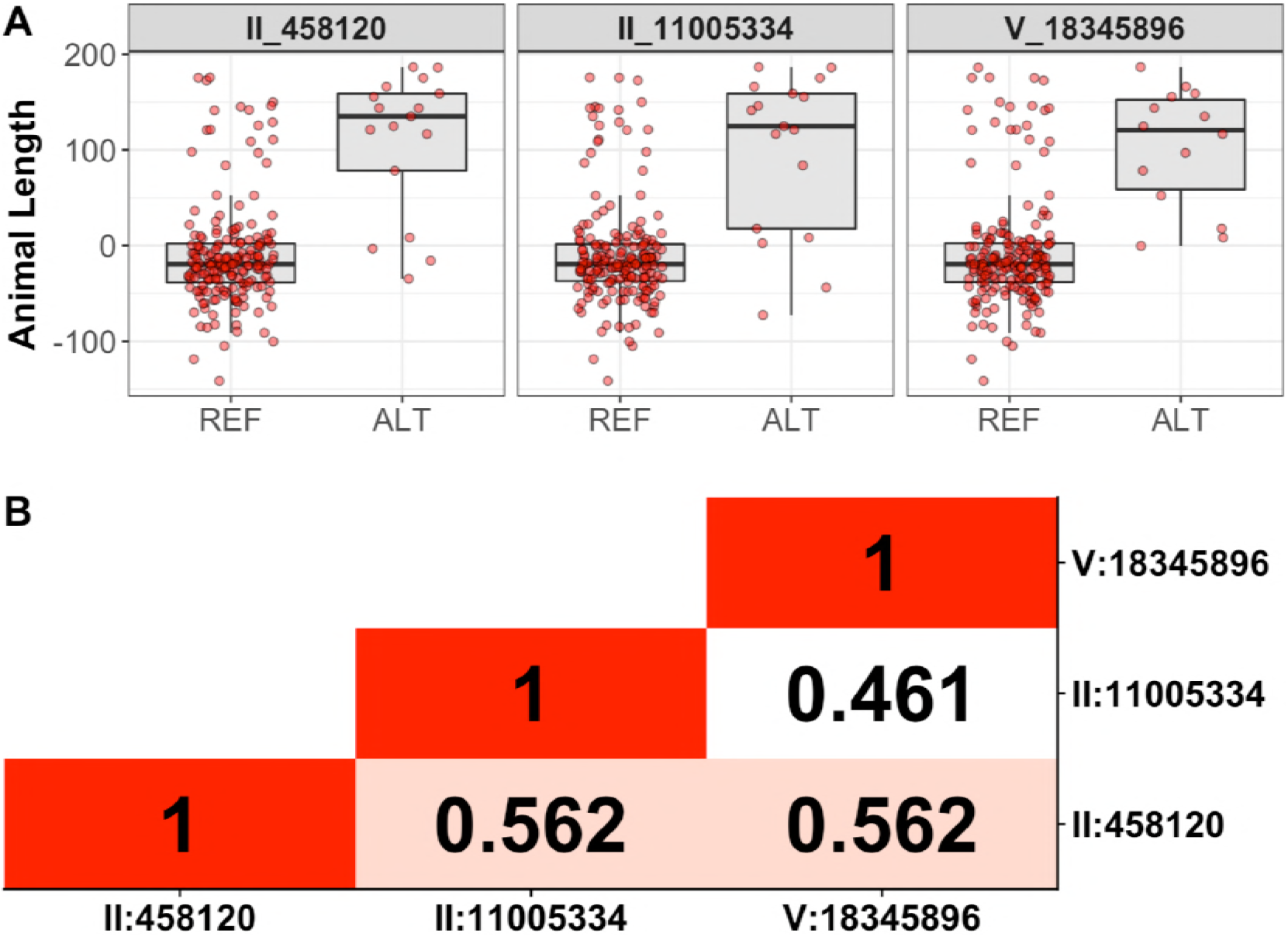
Single-marker mapping summary. A) Regressed animal length (q90.TOF) phenotypes in the presence of ABZ. Each dot represents the mean regressed animal length of four replicates. Strains are grouped by the presence of the REF or ALT genotype at the peak QTL marker identified in the single-marker GWA mapping approach. B) Linkage disequilibrium (LD) as measured by the correlation coefficient between peak QTL markers in A. The formula for the correlation coefficient *r = −D / sqrt(p(A) * p(a) * p(B) * p(b))*.

**Supplemental Figure 3:**
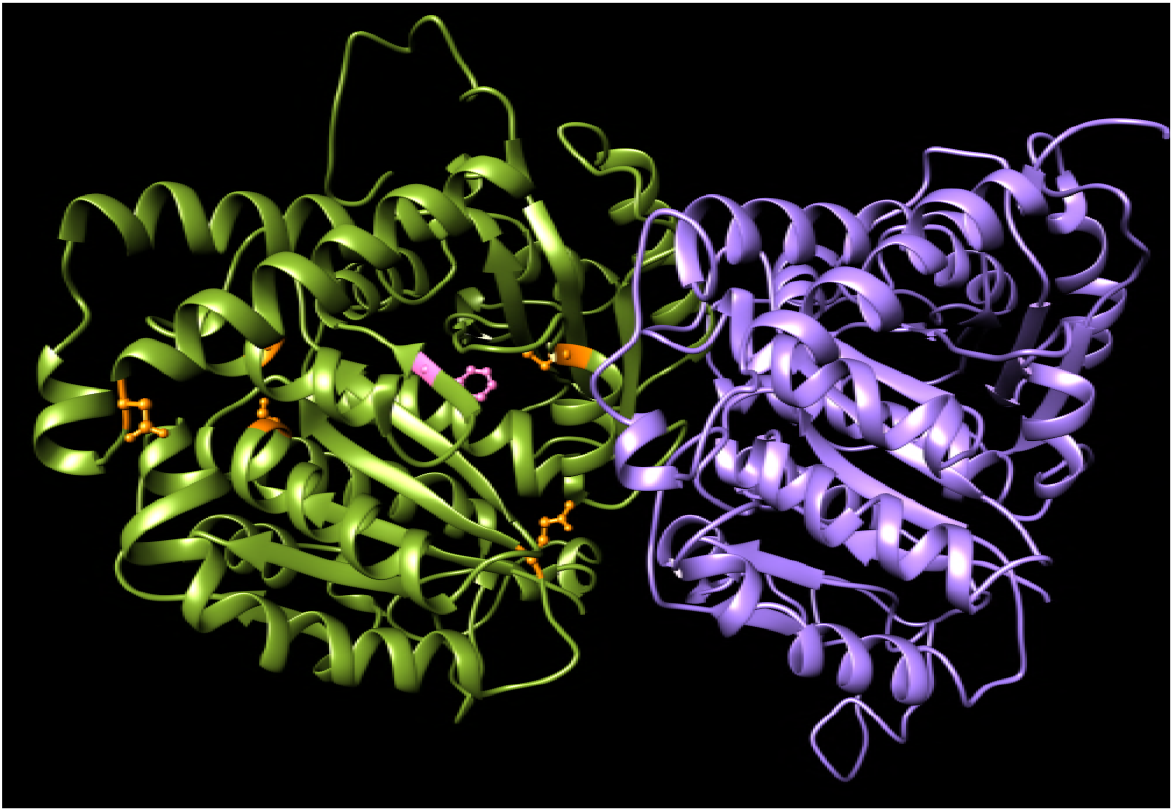
Amino acid substitutions found in *C. elegans* wild strains are distributed throughout the BEN-1 structure. An *in silico* model of the BEN-1 structure (green), binding to an alpha-tubulin (purple) is shown. Novel identified amino acid substitutions among *C. elegans* wild strains are highlighted in color. Most alleles were correlated with ABZ resistance (orange). The F200Y mutation, known as a major BZ resistance marker in parasitic nematodes is shown in pink.

**Supplemental Figure 4:**
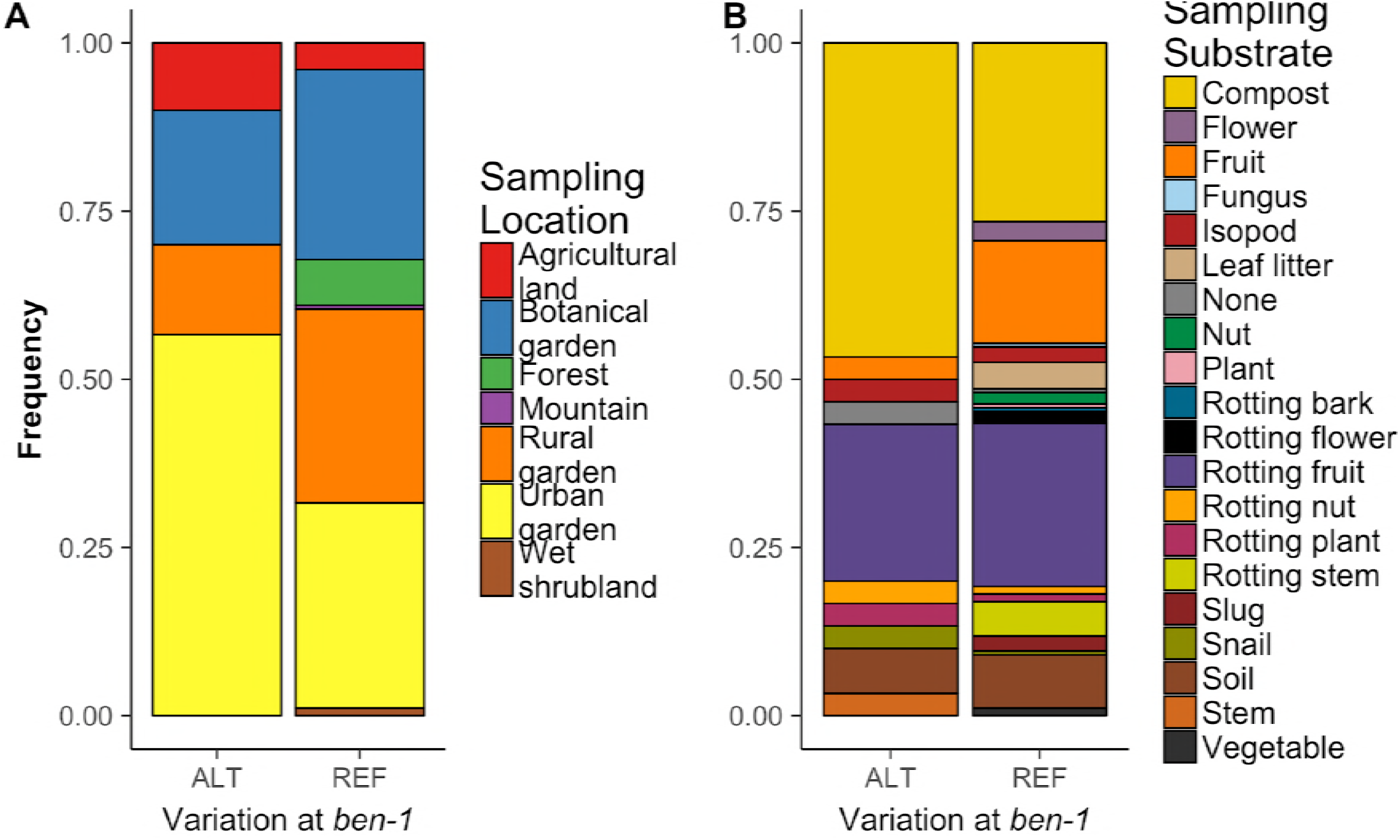
Sampling environment and substrate of wild *C. elegans* isolates. The fractions of wild *C. elegans* isolates sampled in a given A) location and on a given B) substrate are shown. Colors for the stacked bar plots correspond to different A) sampling locations and B) substrates.

**Supplemental Figure 5:**
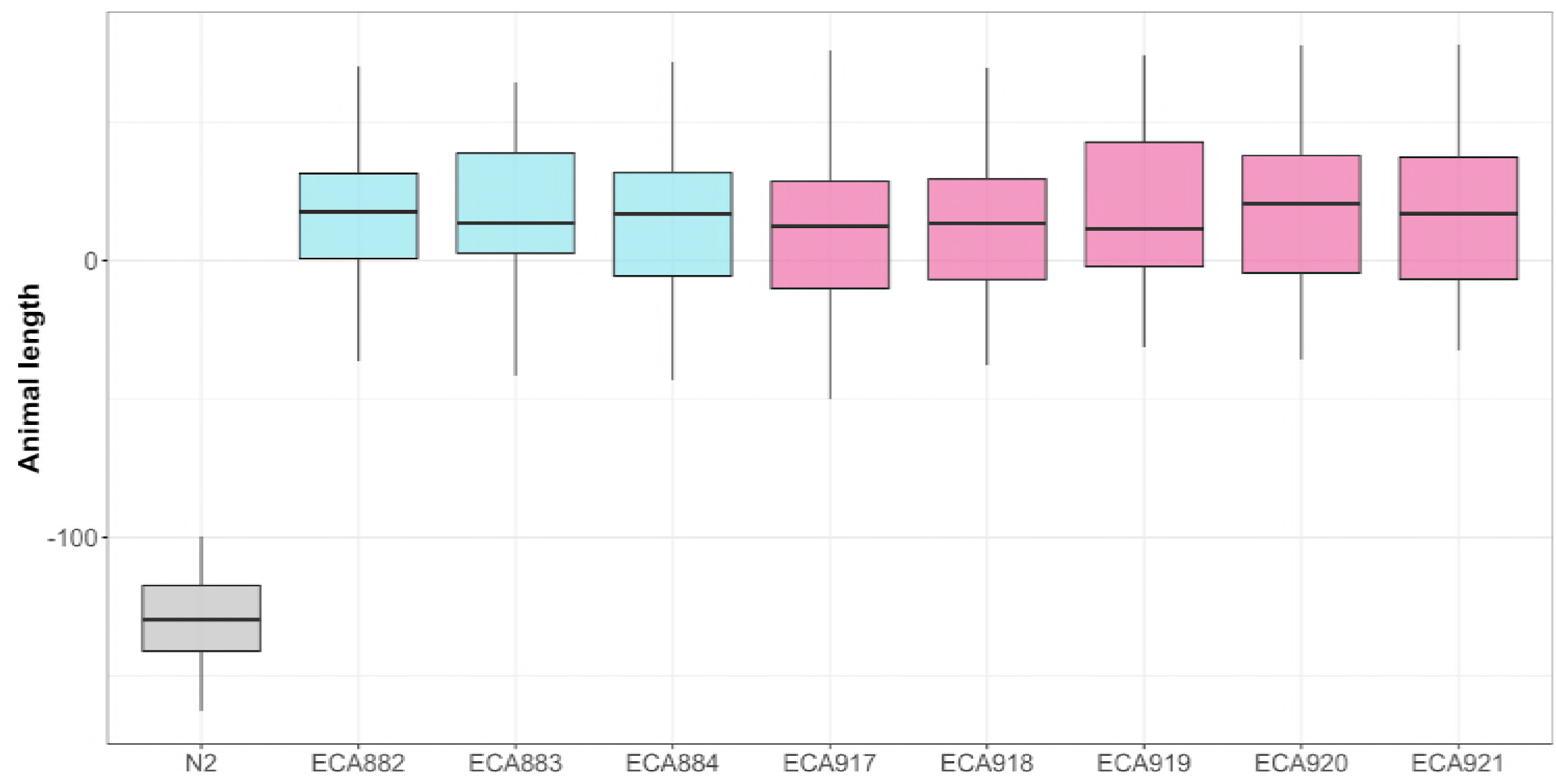
Complete *ben-1* CRISPR allele HTA assay. Tukey box plots of the animal length phenotypes of the generated *ben-1* allele-replacement strains after ABZ treatment is shown. Blue boxes correspond to independent F200Y allele strains and pink boxes correspond to independent Del strains. The y-axis represents the animal-length phenotype after correcting for growth in DMSO conditions.

**Supplemental Figure 6:**
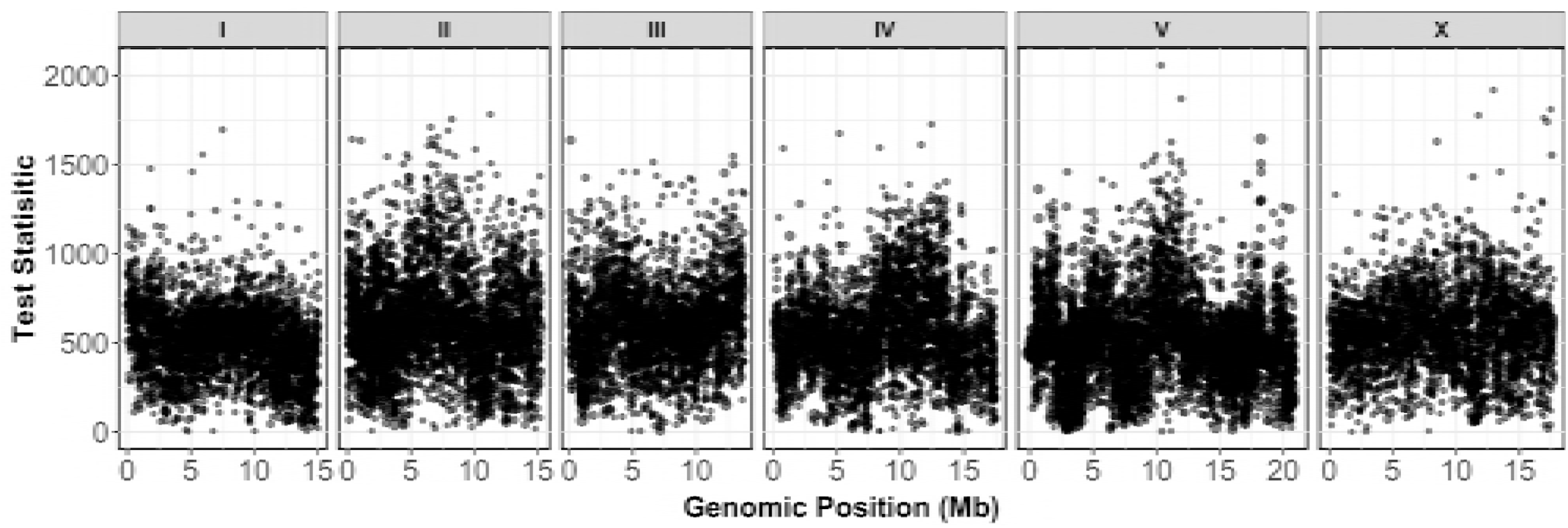
Gene-burden mapping of putative *ben-1* LoF regressed phenotype. Gene-burden manhattan plot of the *ben-1* corrected animal length (q90.TOF) after ABZ exposure. Animal-length phenotypes in the presence of ABZ were adjusted based on the presence of a putative loss-of-function variant in *ben-1*. Each dot represents a single gene of the *C. elegans* genome with its genomic location plotted on the x-axis, and the test statistic plotted on the y-axis. Genes passing the burden test statistical significance threshold are colored in red.

**Supplemental Figure 7:**
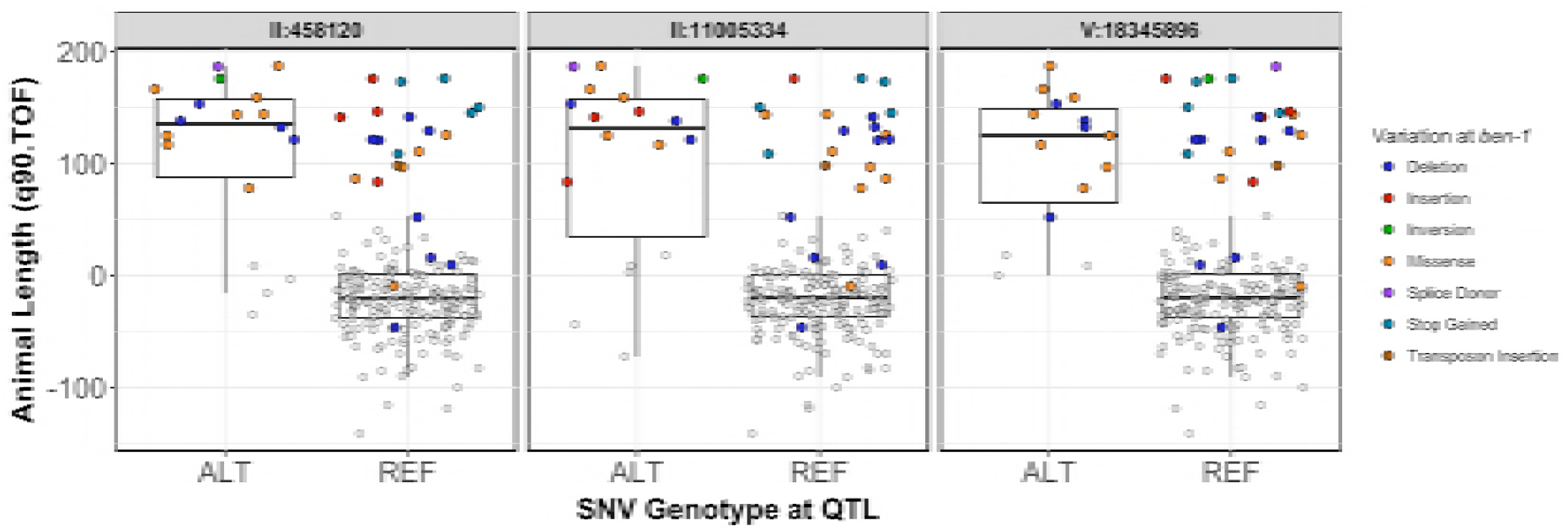
Phenotypes of isolates split by genotypes at peak QTL markers identified using the single-marker mapping approach. Regressed animal length (q90.TOF) phenotypes in the presence of ABZ. Each dot represents the mean regressed animal length of four replicates per strain. Strains are grouped by the presence of the REF or ALT genotype at the peak QTL marker identified in the single-marker GWA mapping approach. Dot colors correspond to the identified variant class in the *ben-1* locus.

**Supplemental Figure 8:**
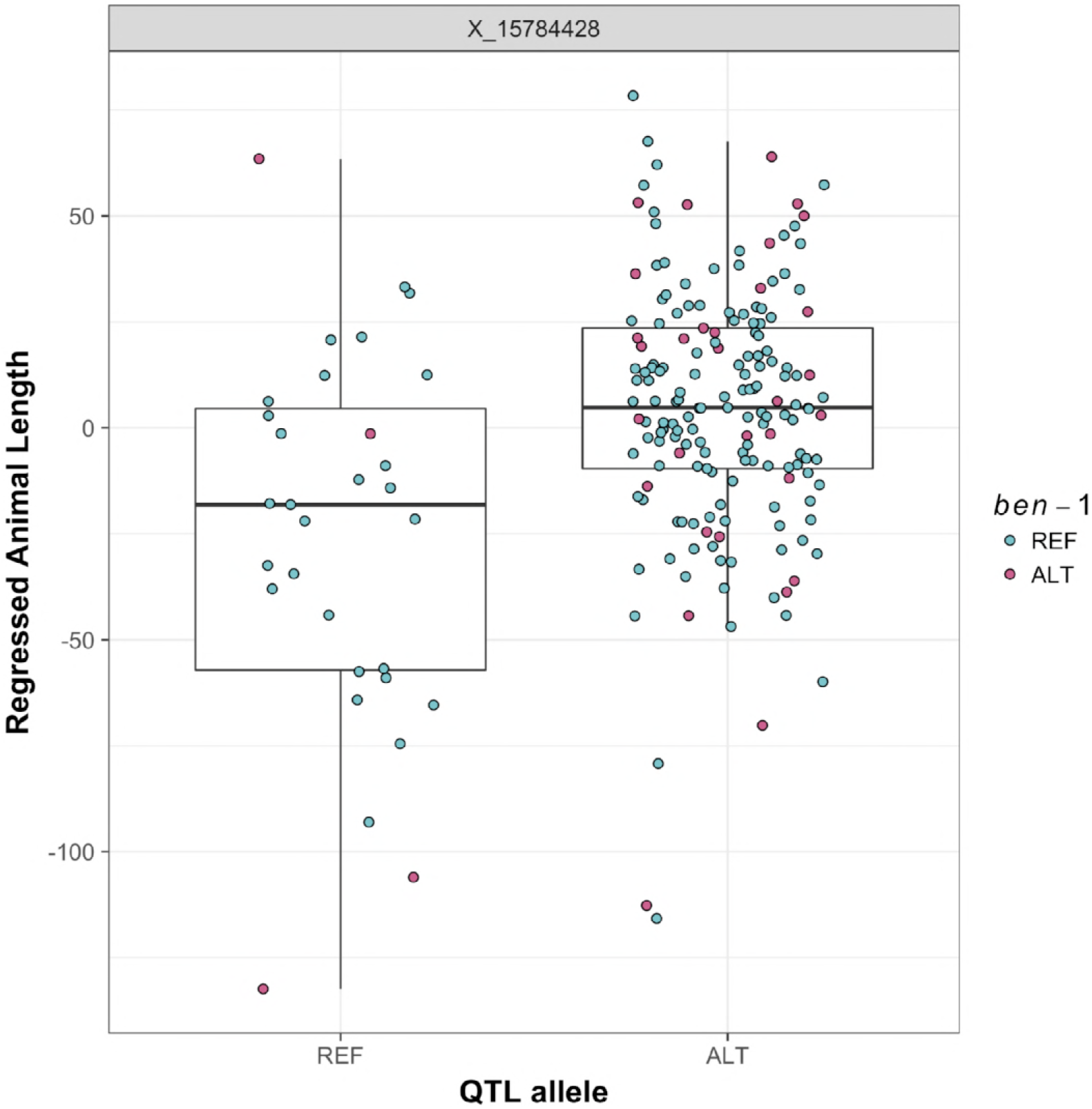
Animal length phenotypes after correcting for the presence of putative *ben-1* loss-of-function variants. Animal length phenotypes in the presence of ABZ after correction for the presence of a putative loss-of-function variant in *ben-1*. Each dot represents the mean regressed animal length of four replicates per strain. Strains are split into groups based on the presence of the REF or ALT allele at the chromosome X QTL peak marker. Dots are colored by the presence of a putative loss-of-function variant in *ben-1*, where pink corresponds to strains with a putative loss-of-function variant, and blue corresponds to strains with no putative loss-of-function variants.

**Supplemental Figure 9:**
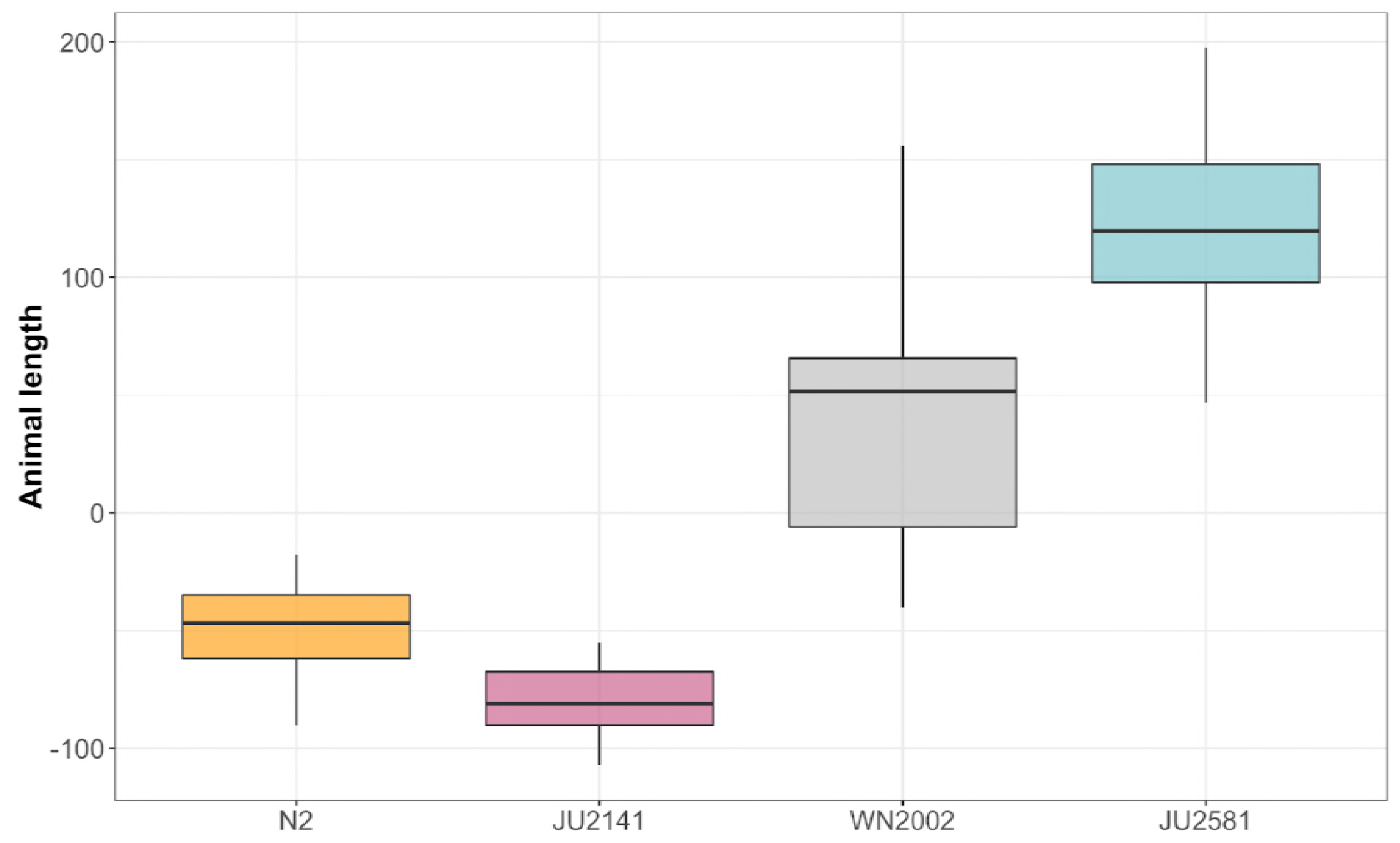
Re-phenotyping of a slow-growing *ben-1* variant strain. Tukey box plots of the animal length phenotypes of N2 and three wild isolates in the presence of ABZ. The y-axis represents the animal length phenotype after correcting for growth in DMSO conditions.

**Supplemental Figure 10:**
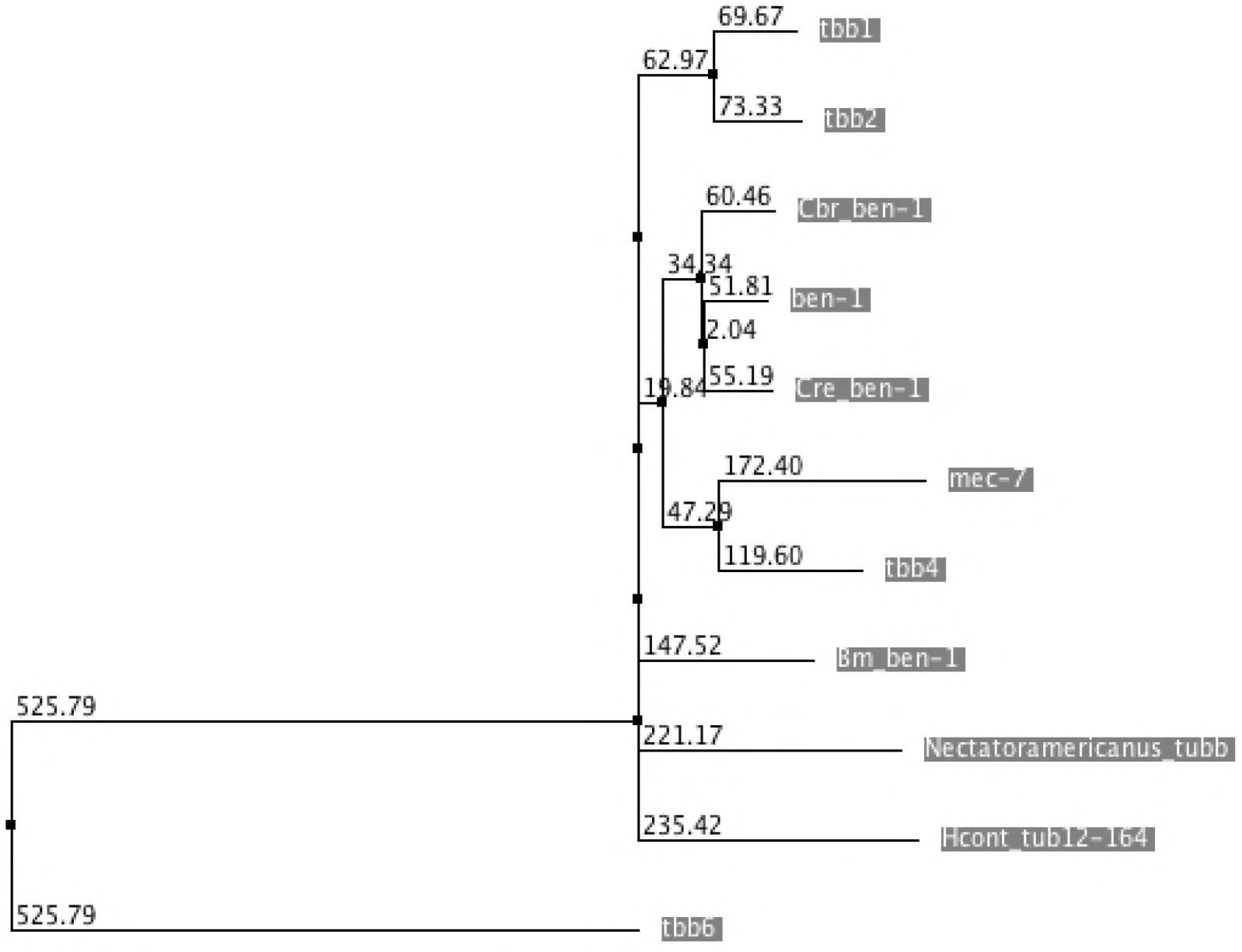
Phylogenetic analysis of β-tubulins. Neighbor-joining tree of the amino acid sequences of the six known *C. elegans* β-tubulins and paralogs in *C. remanei* (Cre), *C. briggsae* (Cbr), *H. contortus* (Hcont), and *Necator americanus*.

**Supplemental Figure 11:**
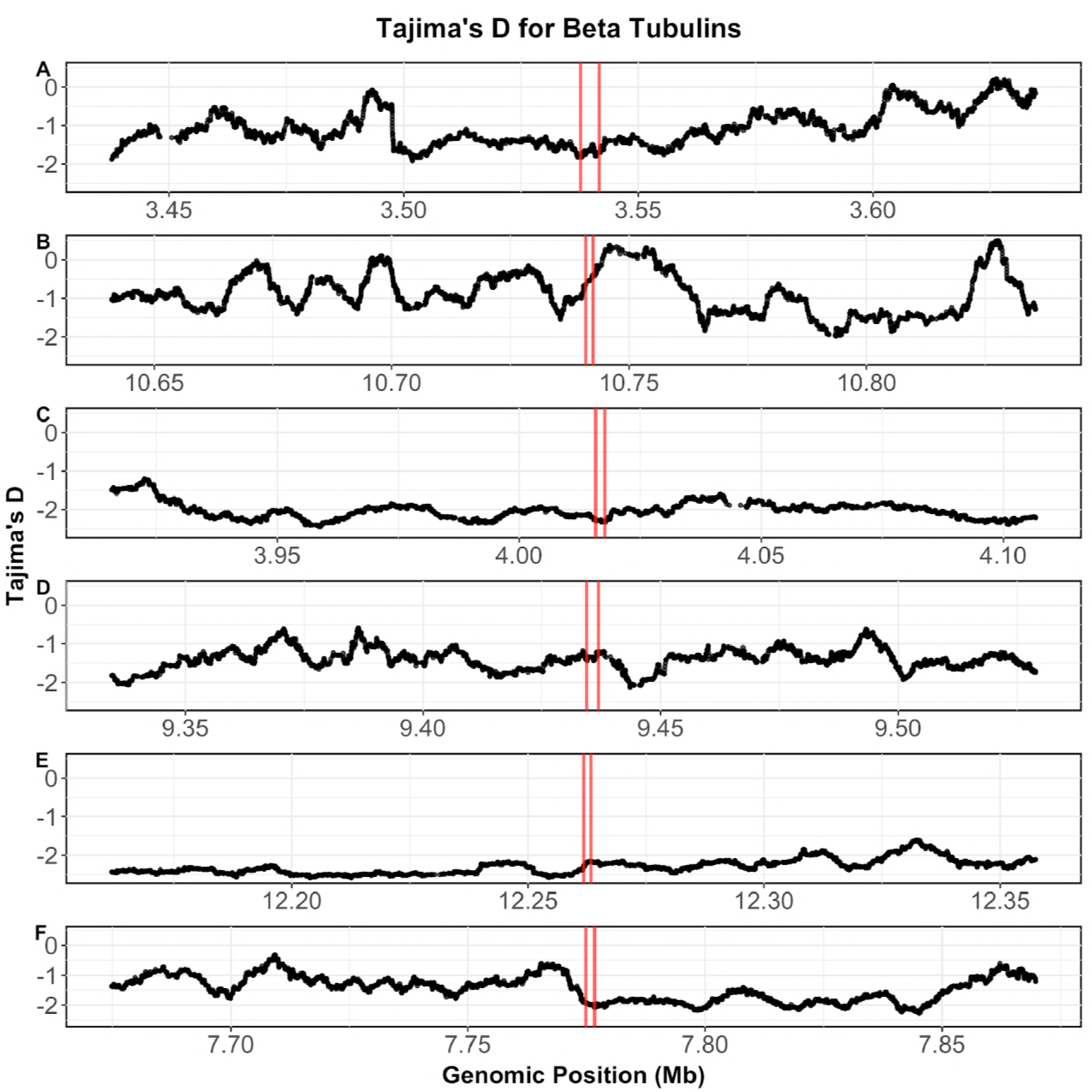
Tajima’s *D* at the *ben-1* locus and other β-tubulin genes. Tajima’s *D* calculated from SNV data of genomic regions surrounding A) *ben-1*, B) *tbb-1*, C) *tbb-2*, D) *tbb-4*, E) *tbb-6*, and F) *mec-7*. Genomic position in Mb is plotted on the x-axis, and Tajima’s *D* is plotted on the y-axis. Tajima’s *D* was calculated using a sliding window with a 100 SNV window and a one-SNV step size. Two red bars for each panel correspond to the start and end positions for the corresponding gene.

